# Projecting the futures of plant traits across habitats in Central Europe

**DOI:** 10.1101/2022.06.06.494936

**Authors:** Marina Golivets, Sonja Knapp, Franz Essl, Bernd Lenzner, Guillaume Latombe, Brian Leung, Ingolf Kühn

**Author notes:** Corresponding author: Marina Golivets, Department of Community Ecology, Helmholtz Centre for Environmental Research – UFZ, Theodor-Lieser-Str. 4, 06120, Halle, Germany.

## Abstract

Many plant traits covary with environmental gradients, reflecting shifts in adaptive strategies under changing conditions and thus providing information about potential consequences of future environmental change for vegetation and ecosystem functioning. Despite extensive efforts to map trait–environment relationships, the evidence remains heterogeneous and often conflicting, partially because of insufficient consideration of distinct trait syndromes for certain growth forms and habitats. Moreover, it is unclear whether traits of non-native and native plant taxa respond similarly to environmental gradients, limiting our ability to assess the consequences of future plant invasions. Here, using comprehensive data for Germany and the Czech Republic and a Bayesian multilevel modeling framework, we assessed relationships between three major plant traits (maximum height, *H*_max_; specific leaf area, *SLA*; and seed mass, *SM*) and environmental factors (7 climate variables and percentage of urban land cover) for native and non-native woody and herbaceous plant assemblages across six broad habitat types. We projected the trait change in these assemblages under future environmental change scenarios until 2081–2100 and quantified the change in trait difference between native and non-native plants. Our models depicted multiple trait–environment relationships, with several important differences attributed to biogeographical status and woodiness within and across habitat types. The overall magnitude of trait change is projected to be greater for non-native than native taxa and to increase under more extreme scenarios. Native woody plant assemblages may generally experience an increase across all three traits, whereas woody non-natives may decline in *H*_max_ and increase in *SLA* and *SM*. Herbaceous *H*_max_ is expected to increase and *SLA* to decrease in most habitats. The obtained trait projections highlight the conditions under which non-native plants may prevail over natives and vice versa and can serve as a starting point for projecting future changes in ecosystem functions and services.

## 1 INTRODUCTION

Economic globalization and human-induced environmental change over the last centuries have caused vast numbers of species to decline (Díaz et al., 2019), while a smaller yet substantial number of species has expanded beyond their historical ranges (i.e., non-native and neonative species; Essl et al., 2019; Seebens et al., 2017). As a result, previously unique species assemblages around the world are increasingly becoming impoverished, more alike, and less stable (Daru et al., 2021; Eichenberg et al., 2021; Finderup Nielsen et al., 2019; Winter et al., 2009; Yang et al., 2021), with serious, often irreversible, consequences for natural ecosystems and humans (Guo et al., 2020; Naeem et al., 2012; Pyšek et al., 2020). In the face of the highly threatened and uncertain future of biodiversity (Thuiller et al., 2005, 2019), it is important to ensure that the scientific knowledge used to design biodiversity policies is easily updatable, synthesizable, and transferable across space and time. This, on the one hand, calls for approaches that generalize scientific outputs beyond individual species and, on the other hand, requires embracing the distinct ecological patterns displayed by different species groups (e.g., native vs. non-native; Liu et al., 2017).

Approaches with a focus on species traits (i.e., any measurable characteristic of a single organism, Violle et al., 2007) are increasingly put forward as a way towards predictive ecology (McGill et al., 2006; Violle et al., 2014) and have been actively employed to study the effects of global environmental change (e.g., Madani et al., 2018; Myers-Smith et al., 2019). The premise of such approaches is that traits mechanistically link an organism’s performance to its environment and can be upscaled to understand and predict how the environment shapes species assemblages and ecosystem functioning (Bjorkman et al., 2018; Dubuis et al., 2013; Küster et al., 2011; Lavorel & Garnier, 2002; Musavi et al., 2016). Moreover, traits yield insights into the mechanisms of non-native species invasiveness (Drenovsky et al., 2012; Küster et al., 2008; Pyšek & Richardson, 2008) and can help reveal differences in the ecological roles and functions of native and non-native species (Hulme & Bernard-Verdier, 2018a, 2018b). However, trait differences between natives and non-natives have mostly been assessed independently of the environmental contexts (e.g., Divíšek et al., 2018; Mathakutha et al., 2019; van Kleunen et al., 2010). In contrast, studies that compare how native and non-native traits shift along environmental gradients and therefore allow extrapolating trait differences into different environmental conditions remain scarce (Gross et al. 2013; Hanz et al., 2022; Henn et al., 2019; Knapp & Kühn, 2012; Sandel & Low, 2019; Westerband et al., 2020). This highlights a mismatch between the trait-based research on native species, which strongly focuses on environmental filtering and adaptation, and that on non-native species, which often insufficiently considers environmental gradients and thus provides only a limited ability to identify the circumstances under which non-natives functionally diverge from or converge with natives.

Despite extensive recent efforts to map trait–environment relationships, the evidence on these relationships remains heterogeneous and often discordant. This might be partly attributed to insufficient consideration of distinct trait syndromes specific to different growth forms and habitats. Notably, woody and herbaceous plants occupy distinct sections in the global spectrum of plant form and function (Díaz et al., 2016), which highlights their unique adaptations to the environment and hence divergent trait–environment relationships (Šímová et al., 2018). Additionally, traits of woody species tend to be more strongly associated with climate than those of herbaceous species (Šímová et al., 2018). This suggests that when all growth forms in a study area are jointly analyzed, trait–environment relationships may appear weak. Nevertheless, it is a common practice for macroecological analyses to pool trait data for woody and herbaceous species together (e.g., Boonman et al., 2020; Moles et al., 2009, 2014; Ordonez et al., 2009; Wright et al., 2005) or to focus only on woody taxa (e.g., Šímová et al., 2015; Swenson et al., 2012), thus hindering generalizations of trait–environment relationships. Moreover, the strength and direction of the associations between traits and environment may vary across different environmental conditions such as represented by habitat types. Most frequently, however, trait–environment relationships have been quantified either as pooled across habitats or for a single specific habitat type per study (e.g., montane open habitats, Dubuis et al., 2013; forests, Maes et al., 2020; Wieczynski et al., 2019). For non-native species, habitat information has been primarily incorporated to compare the levels of invasion across broadly defined (Chytrý et al., 2008) and selected narrowly defined (e.g., grasslands, Axmanová et al., 2021; coastal dunes, Giulio et al., 2020; forests, Wagner et al., 2017) habitats, whereas how traits of non-native species arrange along environmental gradients within or across habitats has not been explored. Collectively, this calls for an explicit consideration of woodiness, habitat type, as well as biogeographical status in trait-based analyses.

In this study, we used extensive plant distribution and trait data for Germany and the Czech Republic and a full Bayesian multilevel modeling framework to assess future trait change following the “assemble first, predict later” approach (Ferrier & Guisan, 2006). We (1) quantified relationships of traits central to plant life history (Díaz et al., 2016; Westoby, 1998) – namely, maximum height, specific leaf area, and seed mass – with climate and land use within native and non-native plant assemblages. Based on obtained trait– environment relationships, we (2) determined the magnitude and direction of plausible future change in mean trait values, which reflect the turnover of taxa and associated functions. The trait change was projected under seven combined climate and socio-economic scenarios for Europe for the period of 2081–2100. Considering the substantial variation in plant adaptive strategies across growth forms and environments, which may weaken or bias the environmental filtering signal if not accounted for (Catford et al., 2021), we addressed each of the goals separately for woody and herbaceous plants and for species pools of six different broad habitat types.

## 2 MATERIAL AND METHODS

### 2.1 Study area

Our study area comprises the entire territory of two Central European countries, Germany and the Czech Republic. Both countries are characterized by a temperate climate with marked regional differences. Mean annual temperature (MAT) ranges from *c.* 4 °C at high elevations to 11 °C in the lowlands, being on average *c.* 8 °C. Total annual precipitation (TAP) averages at 700 mm, ranging from 450 mm in north-eastern lowlands and the east to 1200 mm in the south (Alps) and west (DWD, 2017). Land cover composition across the study area is represented by arable land (37–38%; as of 2018), forested land (30–35%), pastures and mosaic farmland (18–20%), artificial surfaces (7–9%), and semi-natural non-forested land including wetlands and water bodies (1– 4%) (EEA, 2021). The sprawl of artificial surface areas has been the main driver of land-use change over the past two decades in Germany, whereas in the Czech Republic land-use change has been primarily marked by the expansion of forested land (EEA, 2021). For Germany, an increase in MAT by 2.8–5.2 °C and either a decrease or increase in TAP by up to 26% in 2071–2100 (compared to 1971–2000) is projected under severe climate change (RCP8.5) (DWD, 2018). For the Czech Republic, MAT is projected to increase by 4.1 °C and TAP – by up to 16% in 2081–2100 (relative to 1981–2010) under RCP8.5 (Český hydrometeorologický ústav, 2019). Climate change is expected to lead to more frequent extreme drought and rainfall events (Huang et al., 2015; Rulfová et al., 2017; Štěpánek et al., 2016) and increased heatwave impacts (e.g., Krkoška Lorencová et al., 2018) in the study area, among other effects.

### 2.2 Data

#### Taxon-level data

Using multiple open data sources, we collated data on plant taxon occurrences, biogeographical status, habitat affinity, and traits for the entire flora of the study area. We excluded aquatic (i.e., taxa with the Ellenberg moisture indicator value >9), holoparasitic, and fully mycotrophic taxa from analyses.

#### Occurrence records

For Germany, we obtained grid-based native and alien plant taxon occurrence data from the FlorKart database (Datenbank FlorKart, NetPhyD & BfN, 2013) via the information online system FloraWeb (www.floraweb.de; accessed 5 February 2022). FlorKart is the most comprehensive database on plant taxon distribution for Germany, being the result of the combined effort of thousands of volunteers, who were involved in floristic mapping and literature review. From the FlorKart data, which provided the status of each occurrence record, we excluded records of cultivated, erroneous, and doubtful occurrences. For the Czech Republic, we obtained grid-based taxon presence data (i.e., a single record per taxon per grid cell) from the Pladias – Database of the Czech Flora and Vegetation (www.pladias.cz; accessed 16 January 2022). The database resulted from the integration of over 13 million records of *c.* 5,000 species, which originated from multiple national and regional projects, as well as additional data collection efforts within the Pladias project; it is the most complete database on vascular plant occurrence in the Czech Republic (Chytrý et al., 2021; Wild et al., 2019). From the Pladias data, we initially excluded records at the genus level. Records of presently missing or extinct taxa were also removed. Additionally, we excluded data for a few taxa that were known to occur in both countries but for which data were available from a single country only.

The FlorKart and Pladias data were provided at the resolution of the 10’ longitude × 6’ latitude grid cells (corresponding to *c.* 12.0 km × 11.1 km on the 50^th^ parallel). Grid cells in both datasets were originally defined by the sheets (tiles) on the German topographical map (MesstischblalJtter) with a scale of 1:25000 (TK 25), which is commonly used for floristic mapping. Because both databases are essentially compilations of many regional projects with different sampling intensities, we aggregated all data at the grid-cell level (*N* = 3,569) to achieve a more homogeneous sampling effort across the study area. We also cropped the spatial grid to the combined borders of Germany and the Czech Republic and excluded all grid cells with a land area <117 km^2^, which corresponds to the size of the smallest grid cell not truncated by borders or coastlines. To further control for the differences in sampling effort, we excluded grid cells containing less than 83 of the 87 benchmark taxa. Benchmark taxa were taxa that based on the Beals smoothing method (Beals, 1984; Carmona & Pärtel, 2021) occurred in each grid with a probability >0.98. These taxa were determined after all taxonomic names were standardized (see below) and taxa that did not meet the selection criteria (e.g., casual non-natives; see below) were removed. The final grid comprised 3,031 cells, of which 2,481 were located majorly (i.e., >50% of grid cell area) in Germany and 550 in the Czech Republic. As grid cells were delineated by geographic minutes, their size varied with latitude from 117 (in the north) to 139.7 km^2^ (in the south).

#### Biogeographical status

For Germany, we mostly relied on the information on the origin (native, non-native), introduction time (archaeophyte, i.e., introduced before the discovery of the Americas in 1492; neophyte, i.e., introduced after the discovery of the Americas), and establishment status (established, casual, cultivated) from the BiolFlor database (Kühn et al., 2004) and FloraWeb (www.floraweb.de). In case of discrepancies or missing information, we also checked other sources (e.g., Flora Germanica, www.flora-germanica.de), giving preference to local and/or more recent evidence. For the Czech Republic, we used the status information from Pyšek et al. (2012). Using the information on the taxon status, we filtered plant distribution data. Specifically, for taxa that were native in one country and casual non-native in the other country, we retained only records from where those taxa were native. Likewise, for taxa that were native or established non-native in one country but known solely from cultivation in the other country, we considered only data from where those taxa were established. For non-natives that occurred in both countries but were established in only one of them, we kept all known occurrences if the number of occupied grid cells in each country was >1 and discarded singular casual occurrences at the country level. Although retaining occurrences of taxa that were considered casual in a country meant overestimation of the naturalized secondary range size in some cases, more importantly, it allowed us to ameliorate the differences in expert judgment regarding the degree of taxon establishment. Finally, we retained all native taxa and non-native taxa that had established populations in Germany and the Czech Republic in >1 grid cell across the study region. For consistency reasons, we assigned a single, highest achieved degree of establishment to each taxon (i.e., a taxon native in at least one country was treated as native across the study area), which is in accordance with the national treatment of taxa having several statuses in the country.

#### Habitat affinity

To enable analyses at the habitat level, we assigned each taxon to at least one of the following six broad habitat types: forest, heathland and scrub, grassland, wetland, rock and scree, and human-made. Information on taxon habitat affinity was collated from multiple reference sources (namely, BiolFlor, Kühn et al., 2004; Bundesamt für Naturschutz, 2017; EUNIS, Chytrý et al., 2020; DAISIE, Roy et al., 2020; Divíšek et al., 2018; KORINA, www.korina.info, accessed 4 August 2021; Sádlo et al., 2007), which used different habitat classification schemes. We grouped the habitat types in each data source into the six habitat types using expert knowledge (see Table S1 for habitat classification) and then merged all the data. For taxa represented in multiple sources, we only retained habitats listed in the majority of sources, to avoid reports of sporadic occurrences.

#### Traits

We selected three traits for our analyses: (1) typical maximum plant height (*H_max_*; measured in m), (2) seed mass (*SM*; g), and (3) specific leaf area (*SLA*; mm^2^ mg^-1^). These traits depict major plant life strategies (Díaz et al., 2016; Westoby, 1998), correlate with many other important traits (Moles et al., 2007, 2009; Wright et al., 2004), act as both response and effect traits (Hanisch et al., 2020; Kühn et al., 2021; Pollock et al., 2012), and are well represented in open source trait databases (e.g., Kattge et al., 2020). We compiled trait data from multiple databases and online resources: LEDA (Kleyer et al., 2008), TRY (Kattge et al., 2011, 2020, accessed 1 October 2019; see Appendix S2 for references within TRY), EcoFlora (Fitter & Peat, 1994, www.ecoflora.co.uk, accessed 16 September 2021), Info Flora (www.infoflora.ch), iFlora (www.i-flora.com), Kaplan et al., 2019, Vojtkó et al. (2020), World Species (worldspecies.org). We included *H_max_* measurements on vegetative and generative organs, *SM* measurements on dried seeds, and *SLA* measurements on sun and shade leaves and dry biomass. Where possible, we excluded trait measurements from biomes outside our study area (e.g., tundra). Climbers were excluded from analyses of *H_max_*. In most cases, more than one trait value was available per taxon. We used these values to calculate the geometric mean, which is less sensitive to extreme values than other measures of central tendency, after accounting for possible outliers; we calculated *H_max_* as the geometric mean of the maximum height values provided in each data source. Additionally, we categorized each taxon as woody or herbaceous. Woody taxa were defined as perennials whose stems were either entirely lignified or had a lignified base. The information on woodiness was obtained directly or inferred from life form and growth form using the following data sources: Zanne et al., 2014 via the R package *growthform* (v.0.2.3; Taseski et al., 2019); LEDA (Kleyer et al., 2008); BiolFlor (Kühn et al., 2004), Info Flora (www.infoflora.ch), Pladias (www.pladias.cz); TRY (Kattge et al., 2011, 2020); Encyclopedia of Life (eol.org). When the sources provided contrasting information, we assigned woodiness based on morphological descriptions provided in floras. Woody plants occurring in grasslands, wetlands, and rock and scree habitats were excluded from all analyses. Due to the low sample size and variation in trait values, we removed woody archaeophytes.

#### Taxonomic names

To facilitate name harmonization, we initially extracted all the provided names including synonyms for each taxon in FloraWeb and Pladias and matched taxa using those names. This resolved many cases that would be problematic to standardize otherwise (e.g., a hybrid name in one dataset and a corresponding hybrid formula in the other). Subsequently, we checked and updated full taxonomic names against the Plants of the World Online (POWO, 2022) using the R package *taxize* (Chamberlain & Szocs, 2013), Leipzig Catalogue of Vascular Plants using the R package *lcvplants* (v.2.0; Freiberg et al., 2020) and the GermanSL v1.5 checklist using the R package *vegdata* (v.0.9.8; Jansen & Dengler, 2008, 2010). The use of multiple taxonomic databases was necessary because our initial taxon list included many names with qualifiers, hybrid formulas, aggregates, etc., for which none of the available reference sources provided a single optimal solution. Taxonomically critical taxa (e.g., those in genus *Taraxacum*, *Rubus*) were aggregated at higher taxonomic levels (e.g., aggregate, section). Infraspecific taxa were generally aggregated to species or higher levels unless they were non-native.

Our final dataset comprised 1,812 native, 181 archaeophyte, and 331 neophyte taxa; *H_max_* was available for 96%, *SLA* for 74%, and *SM* for 88% of those taxa.

### Climate and land use data

#### Baseline data

We retrieved baseline data on 14 macroclimatic variables from the 10’ × 10’ (*c.* 13 km × 18 km on the 50^th^ parallel) CRU 1961–1990 dataset (New et al., 2002). The variables were total annual precipitation (TAP; mm), precipitation of the driest and wettest quarters (mm), precipitation of the driest month (*P_dry_*; mm), precipitation of the wettest month (*P_wet_*; mm), precipitation seasonality (coefficient of variation of monthly total precipitation, *P_CV_*; %), mean annual temperature (MAT; °C), mean and minimum temperature of the coldest month (°C), mean temperature of the warmest month (°C), maximum temperature of the warmest month (*T_warm_*; °C), mean temperature of the driest quarter (*T_dry_*; °C), mean temperature of the wettest quarter (*T_wet_*; °C), and temperature seasonality (coefficient of variation of monthly average temperature; %). These variables are commonly used in macroecological trait-based studies (e.g., Boonman et al., 2020; Šímová et al., 2018; Wieczynski et al., 2019) and have been projected under different climate scenarios (see below). We rescaled all variables to the 10’ × 6’ spatial resolution by resampling original values onto a 0.5’ × 0.5’ grid and then averaging obtained downscaled values within each 10’ × 6’ grid cell using the R package *raster* (v.3.4-13; Hijmans, 2021). As a baseline for land use, we used Corine Land Cover (CLC) data for the year 2000 (CLC, 2020), which we aggregated to the 10’ × 6’ spatial resolution using the R package *raster* (Hijmans, 2021). Additionally, we determined which of the six habitat types (see Habitat affinity) were present in each grid cell using the Ecosystem types of Europe 2012 raster dataset (EEA, 2018). Forests, grasslands, and human-made habitats were present in all 3,031 grid cells, whereas heaths and scrub occurred in 2,091, wetlands in 966, and rock and scree habitats in 337 grid cells.

#### Scenario projections

We obtained climate and land-use projections for the period of 2081–2100 from the IMPRESSIONS project (www.impressions-project.eu). The core of the project is the IMPRESSIONS Integrated Assessment Platform (IAP2), which combines a suite of sectoral models within a web-based platform and generates quantitative future projections for multiple indicators across Europe at 10’ × 10’ spatial resolution. The IAP2 includes three Representative Concentration Pathways (RCP2.6, RCP4.5, and RCP8.5) and four European Shared Socio-Economic Pathways (Eur-SSP1, Eur-SSP3, Eur-SSP4, and Eur-SSP5; Kok et al., 2019) and permits modeling individual and joint impacts of climate and socio-economic change until 2100. As part of its output, the IAP2 provides per-cell area proportions of seven land-use types (arable, intensive grassland, extensive grassland, urban, managed forest, unmanaged woodland, and unmanaged land). We downloaded climate projections directly from the IMPRESSIONS data repository (ensemblesrt3.dmi.dk/data/IMPRESSIONS; accessed 10 August 2020). To simulate land use projections, we ran the IAP2 for baseline conditions and for 7 combinations of climate and socio-economic scenarios, for each using 3 different dynamically downscaled CMIP5 climate models (listed in Table S2). Next, we matched land use classes in CLC and IAP2 and evaluated the agreement at the 10’ × 10’ resolution between the two baseline datasets. The agreement was high only for urban land, whereas other land-use types showed considerable disagreement (Pearson’s *r* = 0.15–0.68), based on which we used only the percentage of urban land (*U_%_*) in further analyses. We did not consider using the baseline IAP2 projections to parametrize our models because of their inferior spatial resolution and accuracy compared to CLC data.

### 2.3 Data analyses

#### Data preparation

We first log_10_-transformed all traits to reduce the skewness of their distributions and the effect of extreme values. Then, separately for woody and herbaceous native, neophyte, and archaeophyte taxa within selected habitats, we averaged each trait at the grid cell level, omitting taxa with missing trait values. This means that when all the 6 habitat types were present in a grid cell, we computed up to 24 mean values per trait per grid cell (after excluding woody archaeophytes across all habitats and all woody taxa in grasslands, wetland, and rock and scree; see above). The obtained trait mean values were used as response variables in statistical models. Trait means based on less than 4 taxa were heuristically excluded from analyses. For analyses, we separately scaled trait means of woody and herbaceous native, neophyte, and archaeophyte plant assemblages at the habitat level to zero mean and unit variance. We did so because we aimed to capture how taxon status, woodiness, and habitat moderated the effects of climate and land use, rather than to quantify their direct effects on traits. Moreover, such scaling allowed us to impose a single spatial autocorrelation structure across habitats and thus quantify all effects within a single model (as opposed to, for example, fitting a separate model for each habitat type).

To reduce the redundancy among the potential environmental predictor variables, we performed variable selection based on the variance inflation factor (VIF) and Pearson’s correlation coefficient (*r*) using the functions *‘vifstep’* and *‘vifcor’*, respectively, in the R package *usdm* (v.1.1-18; Naimi et al., 2014). Out of the initial 15 predictor variables, we retained 8 (7 variables with VIF <10 and one additional with |*r*| <0.7). Namely, we kept *P*_dry_, *P*_wet_, *P*_CV_, *T*_cold_, *T*_warm_, *T*_dry_, *T*_wet_, and *U*_%_. Prior to analyses, we scaled these variables to the mean of zero and unit variance to aid model parametrization and interpretation. The per-cell number of taxa (*N*_taxa_), which we controlled for in analyses, was scaled similarly to trait values.

#### Statistical models

We assessed trait– environment relationships using linear multilevel models. All models were parameterized within the full Bayesian framework using the Hamiltonian Monte Carlo sampling algorithm implemented in the modeling software Stan (Carpenter et al., 2017) via the function ‘*brm’* in the R package *brms* (v.2.14.4; Bürkner, 2017). We modeled individual trait per-cell mean values as the function of climatic variables and *U*_%_ (continuous predictors) and taxon biogeographical status, woodiness, and habitat (categorical predictors). More specifically, we developed a suite of slope-only models, in which we included a single continuous predictor and its two- and three-way interactions with taxon status and woodiness that were allowed to vary by habitat (i.e., habitat was modeled as a group-level effect). We excluded the main effects of origin, woodiness, and habitat from the models, as those equaled 0 due to the prior scaling of response variables (see Data preparation). The models allowed us (1) to quantify the extent to which the effects of climate and the level of urbanization differ across native and non-native taxa, herbaceous and woody taxa, as well as different habitats, and (2) to incorporate this potential context-dependency into future projections of spatial trait distributions. To account for residual spatial autocorrelation, we included conditional autocorrelation structure (CAR) with grid cell identifier as a grouping factor in all our models. Additionally, in all our models we controlled for *N_spp_*, by including this metric as another predictor variable because sometimes average trait values were correlated with *N_spp_*. In particular, this correlation was negative for *H_max_*, suggesting that taxon-richer grid cells had on average a higher proportion of shorter taxa. Such a pattern may reflect sampling effort (e.g., smaller plants are more likely to remain undetected; Chen et al., 2013) as well as present a genuine ecological phenomenon (Aarssen et al., 2006). In either case, we chose to control for *N_taxa_* in our models, as otherwise its effect could be incorrectly attributed to the environmental predictors. A model for each environmental predictor can be written as follows:

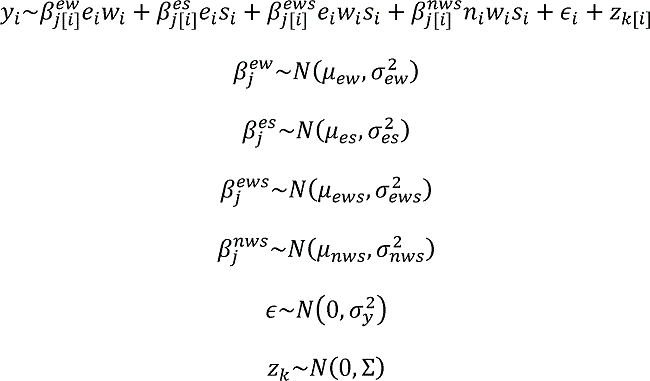

where *y*_i_ is the per-cell mean trait value for the *i*^th^ observation (*i* = 1, …, *N*_obs_), calculated for each combination of habitat type, woodiness and biogeographical status; e_i_ is the environmental predictor; *w*_i_ is woodiness; *s*_i_ is biogeographical status; *n*_i_ is the number of taxa; *β^ew^_j_, β^es^_j_, β^ews^_j_, β^nws^_j_* are slopes for the interactions between variables indicated in the superscript in the *j*^th^ habitat type (*j* = 1, …, *N*_habitats_); *β^es^_j_* is the slope for the interaction between *e*_i_ and *s*_i_ in the *j*^th^ habitat type; *β^ews^_j_* is the slope for the interaction between *e*_i_, *w*_i_ and *s*_i_ in the *j*^th^ habitat type; ∈_i_is the residual effect of the *i*^th^ observation; z_k_ is the residual spatial random error for the *k*^th^ grid cell (*k* = 1, …, *N*_cells_); µ^2^_ew_, µ^2^_es_, µ^2^_ews_, µ^2^_nws_ are the overall slopes for the interactions specified in the subscript; are the habitat-level variances for the slopes; is the residual variance; and Σ is the covariance matrix, as defined in a conditional autoregressive model. We chose to fit separate models for each environmental predictor because of the high complexity of a full model (i.e., for each environmental predictor, we estimated 12 model parameters) and possible collinearity due to a large number of interaction terms with the same categorical predictors.

To prevent the sampler from considering highly implausible values, we used zero-centered weakly informative priors, which we chose based on prior predictive checks (Wesner & Pomeranz, 2021). For each model, we ran 4 chains with 10,000 iterations per chain, starting from default values. We discarded the first half of each chain as a warm-up and thinned the other half at the interval of 4, which resulted in 5000 draws from the posterior distribution for each parameter. The potential scale reduction factor, Ȓ (Gelman & Rubin, 1992), was close to 1 for all our models, indicating convergence.

### Model predictive performance

We evaluated model predictive performance using exact k-fold cross-validation. To avoid the potential overestimation of predictive performance, the folds were determined as spatial blocks (Roberts et al., 2017). For that, we overlaid a 3 × 3 spatial-block grid onto the grid of the study area (Figure S1) using the function *‘spatialBlock’* in the R package *blockCV* (v2.1.4; Valavi et al., 2019), which resulted in 8 spatial blocks, two of which we subsequently merged to achieve a more even distribution of grid cells across folds. We then assigned all data points within a grid cell to a specific fold and performed a 7-fold cross-validation with the function *‘kfold’* in the R package *brms* (Bürkner, 2017). As the measures of model predictive performance, we calculated the k-fold information criterion (*kfoldIC*) and the root mean square error based on cross-validated predictions (*RSME*; Table S3).

### Future projections

For each trait, we obtained projections of per-cell mean trait values on baseline and scenario data (7 RCP–SSP scenario combinations × 3 dynamically downscaled CMIP5 climate models for each RCP; see Table S1). We set the scaled *N*_taxa_ to 0 in all projections. These projections were computed as the weighted average of posterior predictive distributions from the eight models each with a single environmental predictor variable. For averaging, we used Bayesian stacking of predictive distributions (Yao et al., 2018), a method that explicitly optimizes individual model weights to maximize the leave-one-out predictive density. As input for Bayesian stacking, we provided the expected log pointwise predictive densities based on the results of k-fold cross-validation. We then calculated the projected per-cell change in each trait under each scenario as the difference between the medians of the projected future and baseline posterior predictive distributions. To enable comparison across biogeographic statuses, we rescaled trait change values of archaeophytes and neophytes to the standard deviations (SD) of a baseline trait distribution for native taxa of corresponding woodiness and habitat. To assess the overall magnitude of trait change, we calculated a Euclidean distance between projected per-cell posterior means of the three traits on the baseline data and scenario data.

To summarize the effects of all environmental predictors across all three traits and to visualize the direction of those effects in the multivariate space, we performed separate redundancy analyses (RDA) on the subsets of the projected trait change that corresponded to all unique combinations of biogeographical status, woodiness, and habitat across all the scenarios using the R package *vegan* (v.2.5-7; Oksanen et al., 2020). As the measure of individual predictor contribution, we calculated the length of the vectors with the initial point at (0,0) and the terminal point at the scores of the first two RDA axes. The vectors reflected the weighted effect sizes of the predictors that were used to calculate posterior predictive distributions. All statistical analyses and visualizations were performed in the R environment v4.1.0 (R Core Team, 2021).

## 3 RESULTS

Overall, we observed a high degree of variability (i.e., the spread across posterior distributions) and uncertainty (i.e., the spread within posterior distributions) in the magnitude and direction of the projected per-cell change in maximum height (*H*_max_), specific leaf area (*SLA*), and seed mass (*SM*) under environmental change scenarios. The variation in the projected trait change was pronounced at all three grouping levels considered in the analyses, i.e., taxon biogeographical status, woodiness, and habitat type, and increased with the degree of environmental change (Figures 1–4, S2–S7, S14). Likewise, the degree of uncertainty associated with individual per-cell projections varied across the grouping levels and was generally higher under more extreme scenarios, being driven much more by climate change than urbanization (Figures S8–S13). The lowest likelihood of trait change (i.e., >50% grid cells with projected posterior credible intervals excluding zero) was observed under RCP4.5 and RCP8.5 for woody native *SLA* in human-made habitats and for archaeophyte *SM* in human-made habitats and heaths and scrub.

**FIGURE 1.**
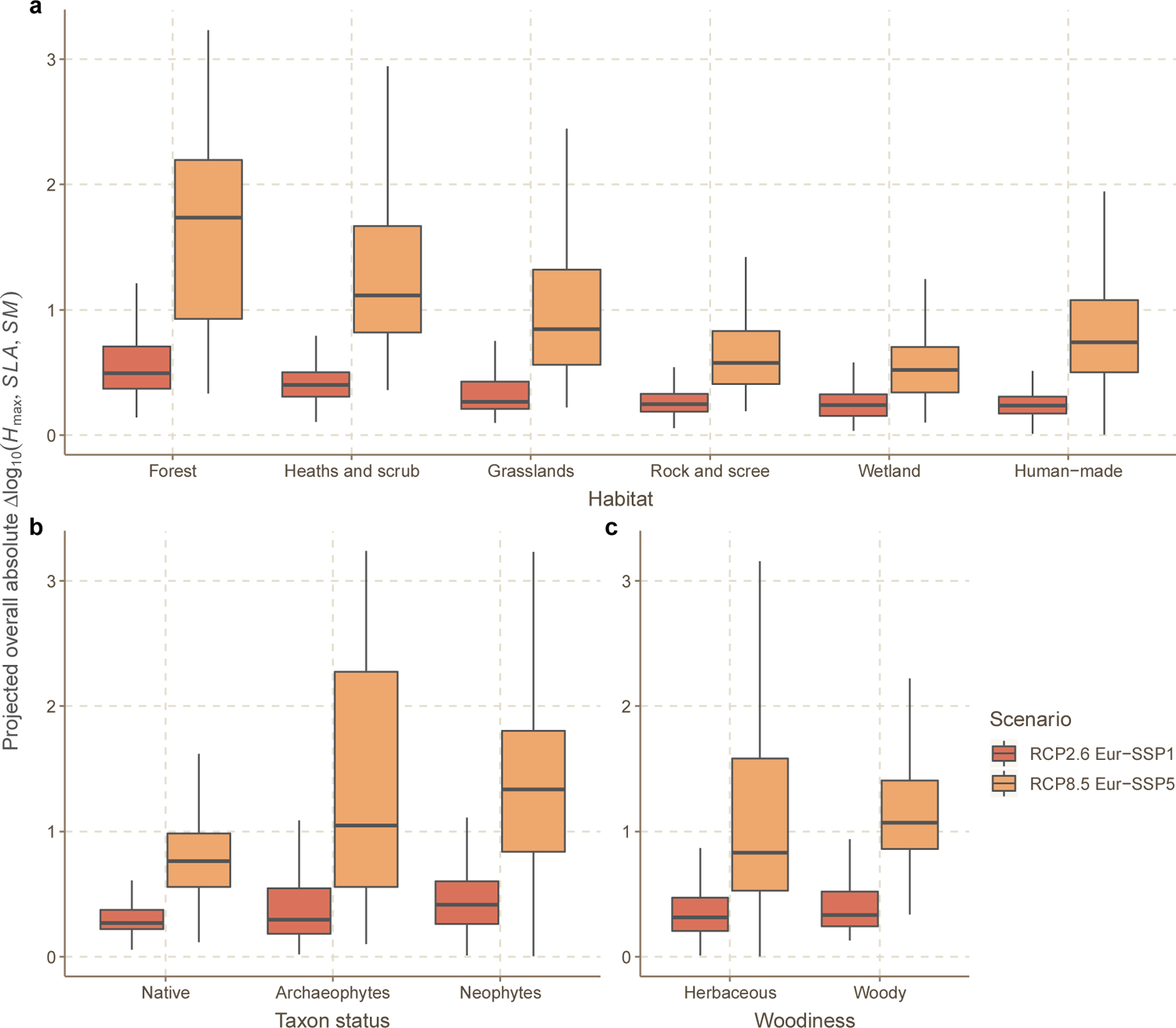
The projected absolute overall per-cell change across the three traits, plant maximum height (*H*_max_), specific leaf area (*SLA*), and seed mass (*SM*), at the habitat (a), biogeographical status (b), and woodiness (c) levels under the least extreme combined climate and socio-economic scenario (RCP2.6 Eur-SSP1, all climate models pooled) and the most extreme one (RCP8.5 Eur-SSP5) for 2081–2100. The overall trait change was calculated as a Euclidean distance between projected per-cell posterior means of the three traits on baseline data and scenario data. Boxes show 25%, 50%, and 75% quartiles, and whiskers show 95% credible intervals.

**FIGURE 2.**
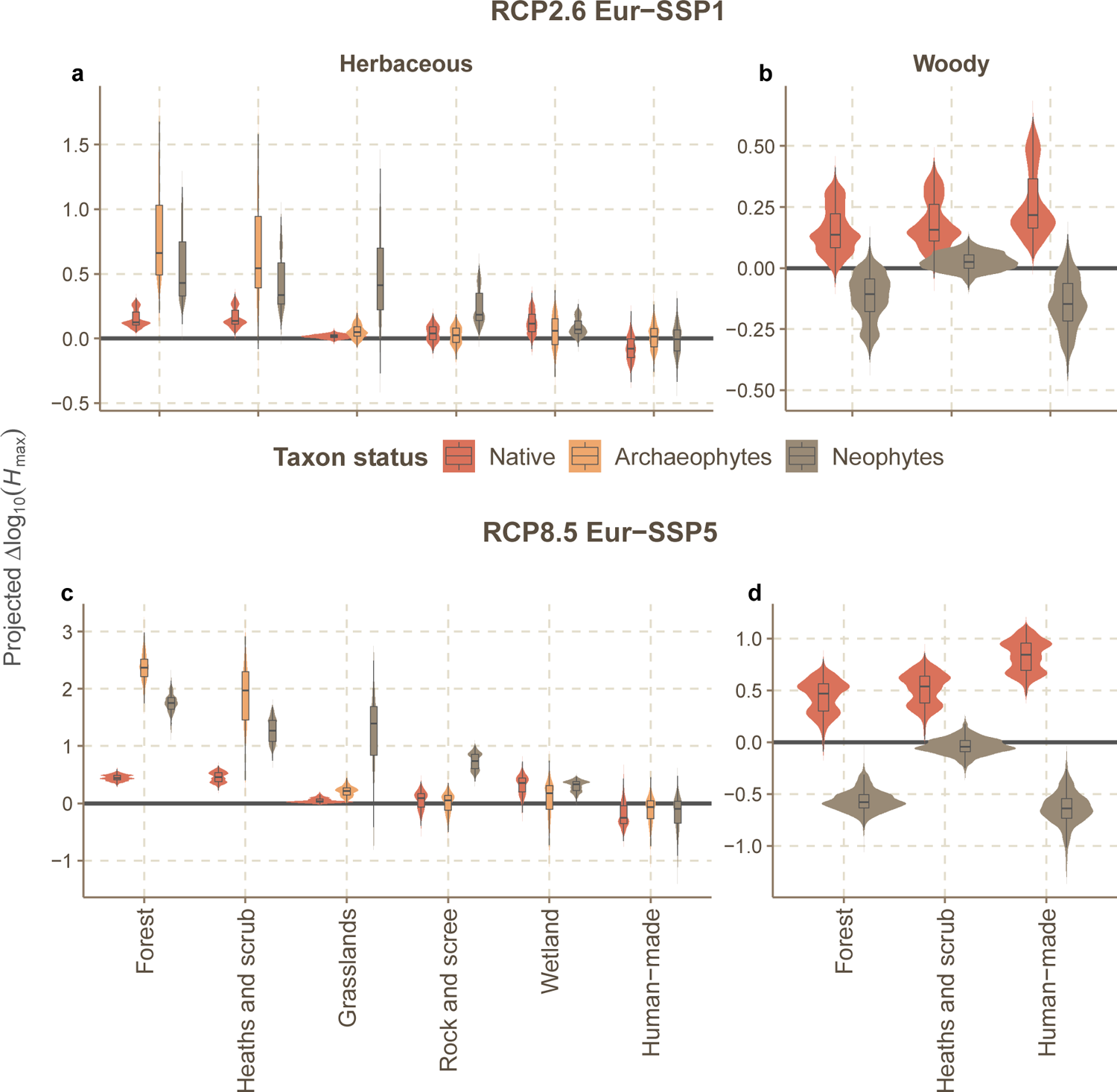
The projected per-cell (10’ × 10’) change in the log_10_-transformed maximum plant height (*H*max) under the least extreme combined climate and socio-economic scenario (RCP2.6 Eur-SSP1, all climate models pooled) and the most extreme one (RCP8.5 Eur-SSP5) for 2081– 2100, for herbaceous (a, c) and woody (b, d) taxa in six broad habitat types. The trait change here is expressed as the posterior means of per-cell model predictions. The violin plots depict the distributions of predicted values across the study area and climate models (Table S1) and the boxplots provide summary statistics of those distributions (boxes show 25%, 50%, and 75% quartiles and whiskers give roughly 95% credible intervals). For each habitat by woodiness combination, the trait change is presented in standard deviations (*SD*) of the baseline trait distribution of native taxa for that combination. For example, the overall change of 0.60 in *H*_max_ of forest herbaceous neophytes under the RCP2.6 Eur-SSP1 scenario indicates that the average *H*_max_ of this assemblage is projected to increase by 0.60 *SD*, relative to the current *H*_max_ distribution of natives. Note the different scaling of Y-axes. Projections under other scenarios are illustrated in Figures S2, S3 (projected per-cell posterior means), S8, S9 (projected per-cell posterior standard deviations).

**FIGURE 3.**
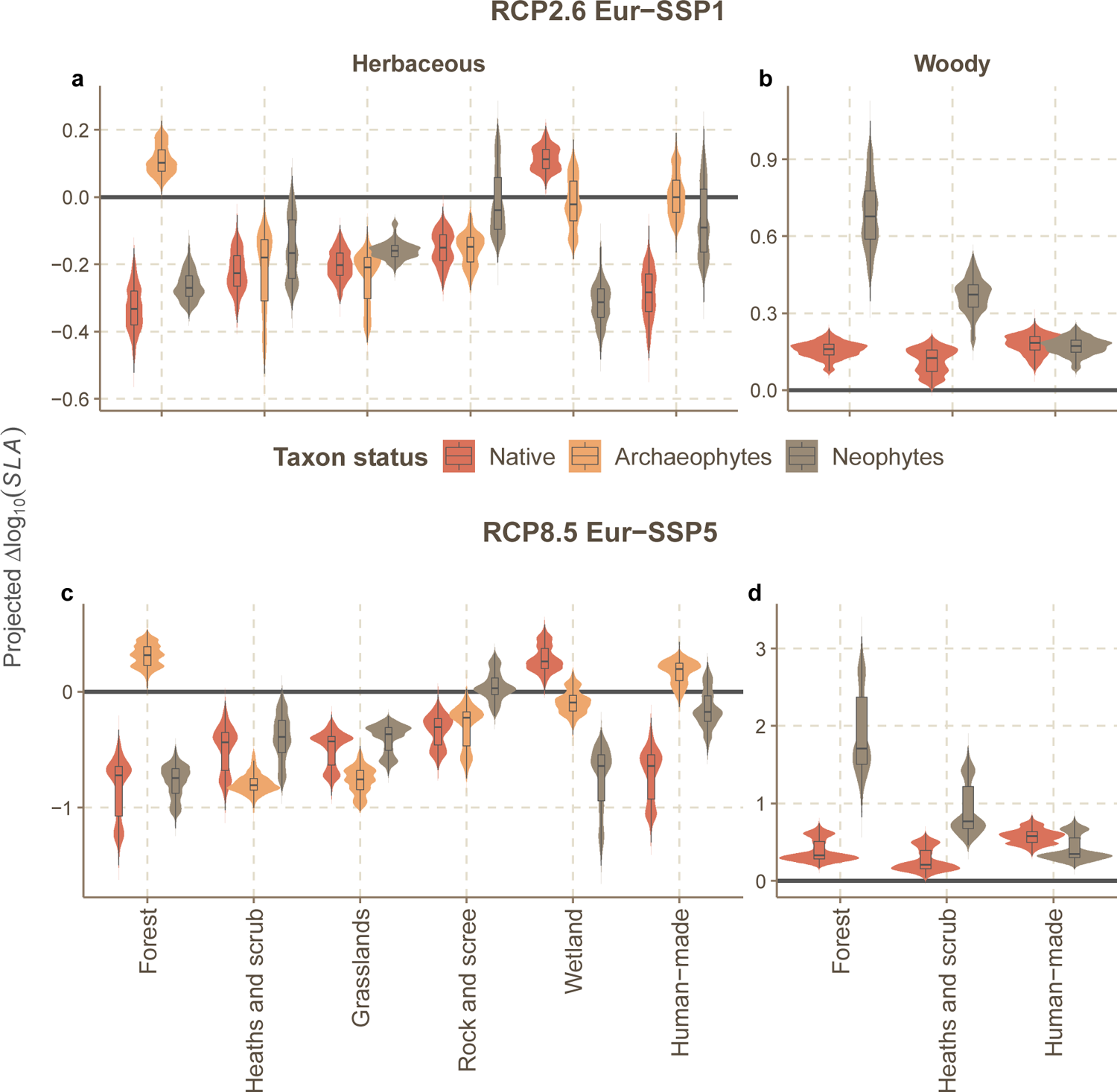
The projected per-cell (10’ × 10’) change in the log_10_-transformed specific leaf area (*SLA*) under the least extreme combined climate and socio-economic scenario (RCP2.6 Eur-SSP1, all climate models pooled) and the most extreme one (RCP8.5 Eur-SSP5) for herbaceous (a, c) and woody (b, d) taxa in six broad habitat types. Projections under other scenarios are illustrated in Figures S4, S5 (projected per-cell posterior means), S10, S11 (projected per-cell posterior standard deviations). Other details are as in Figure 2.

**FIGURE 4.**
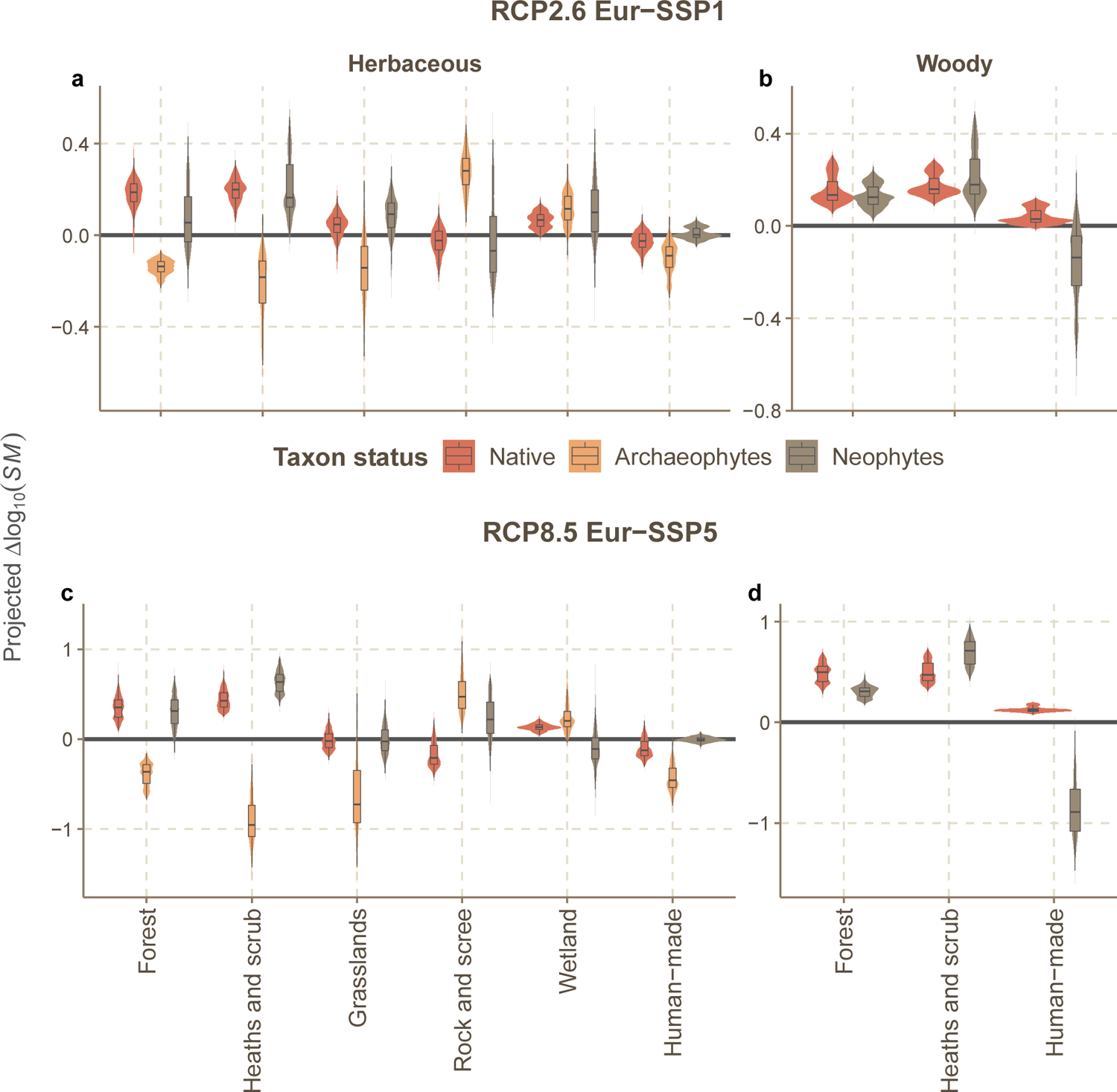
The projected per-cell (10’ × 10’) change in the log_10_-transformed plant seed mass (*SM*) under the least extreme combined climate and socio-economic scenario (RCP2.6 Eur-SSP1, all climate models pooled) and the most extreme one (RCP8.5 Eur-SSP5) for 2081–2100, for herbaceous (a, c) and woody (b, d) taxa in six broad habitat types. Projections under other scenarios are illustrated in Figures S6, S7 (projected per-cell posterior means), S12, and S13 (projected per-cell posterior standard deviations). Other details are as in Figure 2.

Below, we focus on the posterior means of the projected per-cell posterior distributions, which reflect the overall trends in our projections but do not embrace the uncertainty, for the least and most extreme scenarios (RCP2.6 Eur-SSP1 and RCP8.5 Eur-SSP5, respectively) to illustrate the maximum future option space for 2081–2100. The results for other scenarios are presented in the Supplementary Material.

### 3.1 Projected trait change

When all traits were considered, the magnitude of trait change was expected to be on average higher in forests, heaths and scrub, and grasslands than in other habitat types (Figure 1a), for non-natives than natives (Figure 1b), and for woody than herbaceous plants (Figure 1c). The direction of the projected individual trait change often diverged for herbaceous vs. woody plants as well as for natives vs. neophytes (Figure S14).

In all scenarios, the mean trend for *H*_max_ of herbaceous plant assemblages in tree- and shrub-dominated habitats was positive throughout the study area, being more pronounced for archaeophytes and neophytes than for natives (Figures 2a,c, S2a–f). In other habitats, the change of *H*_max_ was expected to be more heterogeneous spatially and across biogeographical statuses. Notably, in grassland and rock and scree habitats, herbaceous native *H*_max_ was projected to show very little to no change, whereas herbaceous neophytes were anticipated to increase in *H*_max_, especially in grasslands (Figure 2a,c, S2g–l, S15a,c). As for woody plant assemblages, only *H*_max_ of natives demonstrated a predominantly positive trend, with the highest increase projected for human-made habitats under RCP8.5 SSP5; meanwhile, *H*_max_ of neophytes tended to mostly decrease (Figures 2b,d, S3, S15b,d).

Similar to *H*_max_, the projected change of *SLA* varied with taxon biogeographical status, woodiness, and habitat (Figures 3, S4–5). Herbaceous *SLA* was projected to mainly decrease, with some exceptions (e.g., native *SLA* in wetlands; Figures 3a,c, S4). At the same time, the degree of this decrease tended to be lower for natives than neophytes (S16a,c). In contrast, woody *SLA* was anticipated to increase for both natives and neophytes, with the latter increasing more than natives in forest and heaths and scrub habitats (Figures 3b,d, S5, S16b,d).

The overall magnitude of *SM* change was comparable to that of *H*_max_ and *SLA*. In forests and heaths and scrub, woody native and non-native *SM* was projected to only increase, whereas in human-made habitats neophyte *SM* showed a decline (Figures 4b,d, S7a–e, S17b,d). The direction of projected change in herbaceous *SM* was less uniform and varied spatially and with habitat and biogeographical status (Figures 4a,c, S6, S17a,c). Particularly, *SM* of archaeophytes tended to respond opposingly to natives and neophytes and was more likely to decrease in the majority of habitats (Figures 4a,c, S6).

### 3.2 Contribution of environmental predictors

Across all three traits, *T*_warm_ and *T*_cold_ captured the highest amount of variation in the projected trait change, followed by *P*_CV_, *P*_dry_, and *P*_wet_; the contributions of *T*_dry_, *T*_wet_, and *U*_%_ were considerably smaller (Figure S18). The role of individual environmental predictors generally varied with taxon biogeographical status, habitat, and woodiness (Figure 5). Nonetheless, some predictors showed highly consistent associations with the projected trait change. For example, a projected increase in herbaceous *SM* correlated mostly positively with *P*_CV_ and negatively with *P*_dry_; the opposite was, however, observed for herbaceous neophyte *SM* in forest, heaths and scrub, and rock and scree habitats. Similarly, *T*_cold_ contributed positively to the *SM* increase in woody plant assemblages, apart from neophytes in human-made habitats. Meanwhile, *T*_warm_ and *T*_cold_ exhibited a strong positive relationship with both herbaceous and woody *H*_max_, except for human-made habitats and woody neophytes, for which the relationship was reverse. In some cases, the effect of temperature on *H*_max_ was not as pronounced as that of *P*_wet_ and *P*_CV_. Particularly, native *H*_max_ in grasslands and human-made habitats was strongly positively affected by *P*_wet_ and *P*_CV_, whereas in rock and scree habitats native *H*_max_ was negatively associated with the two predictors. With few exceptions, herbaceous *SLA* correlated negatively with *T*_warm_, *T*_cold_, and *P*_dry_. Additionally, native herbaceous *SLA* positively correlated with *P*_wet_ in all habitats apart from wetlands, whereas non-native herbaceous *SLA* consistently showed a negative relationship with *P*_wet_. Unlike herbaceous *SLA*, native and neophyte woody *SLA* was strongly positively related to *T*_warm_, *T*_cold_, and *P*_dry_ (Figure 5).

**FIGURE 5.**
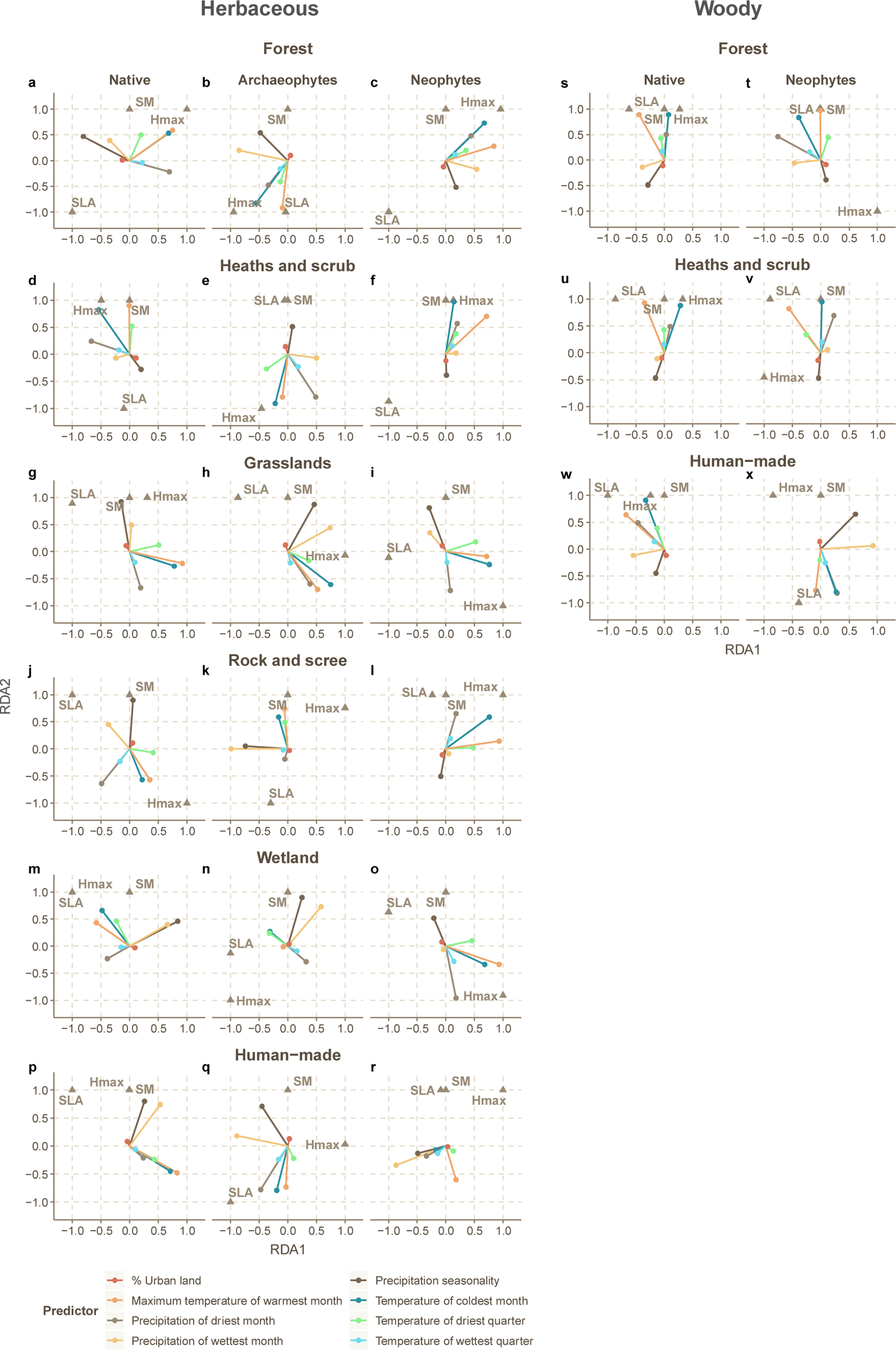
The relative contribution of individual environmental predictors to the projected change in herbaceous (a–r) and woody (s–x) plant maximum height (*H*max), specific leaf area (*SLA*), and seed mass (*SM*), calculated using redundancy analysis (RDA). A separate RDA was performed on each subset (*N* = 24) of per-cell projections across all the scenarios, representing a unique combination of taxon biogeographical status, habitat, and woodiness. Shown are the RDA scores of the predictors (as vectors) and traits (as triangles; centered at seed mass) for the two first RDA axes (RDA1, RDA2). RDA1 and RDA2 together captured 91–100% of the variation in the data. The lengths of vectors are proportional to the magnitude of the effects of respective predictors on the three traits simultaneously. The overall contributions of each predictor, calculated as the combined length of respective vectors across all the RDA spaces, are shown in Figure S18. The angles among the vectors and triangles reflect their correlation, which equals the cosine of the angle. For example, the angle between *SM* and precipitation seasonality is acute for most assemblages, indicating their strong positive correlation.

## 4 DISCUSSION

In this study, we quantified the broad-scale relationships of three traits central to plant life history – maximum plant height (*H*_max_), specific leaf area (*SLA*), and seed mass (*SM*) – with eight selected environmental variables, and used the obtained relationships to project the trait change under seven plausible scenarios of future environmental change in Central Europe. We explicitly modeled the variation in trait–environment relationships associated with plant woodiness, biogeographical status, and habitat type, to account for the fact that different types of plant assemblages may exhibit unique adaptations to the environment, hence making some aspects of mechanistic context-dependence explicit (Catford et al., 2021). We showed that the three traits were projected both to increase and decrease to varying degrees across and – in many cases – within habitats and that the overall magnitude of this change was expected to be on average higher for non-native than native taxa (Figure 1b) and under more extreme scenarios (Figures S2–S7). Moreover, we found that in the future, distinct environmental responses of native and non-native plants may lead to even higher trait values than currently observed for non-natives (e.g., herbaceous *H*_max_ in most habitats) as well as to a reduced average native–non-native trait difference (e.g., woody *H*_max_; Figures S15–17), which may result in altered competitive hierarchies among natives and non-natives (Kunstler et al., 2016).

### 4.1 Projected overall trait change across habitat types

The overall magnitude of trait change was projected to be higher in forests, heaths and scrub, and grassland habitats compared to rock and scree, wetland, and human-made habitats (Figure 1a). This result, at least for wetlands and rock and scree habitats, may reflect both the low sensitivity of these habitat types to the environmental change as well as the limitations of our approach to capture environmentally-driven trait variation at smaller scales. In particular, rock and scree habitats are highly stable systems defined by environmental stress more than by macroclimate, whereas wetlands are strongly shaped by local hydrology, which we did not account for, and show much less turnover along macroclimatic gradients than other habitats. Moreover, the spatial resolution of our analyses might have been too coarse to capture the environmental gradients that traits of wetland and rock and scree plant assemblages respond to. Human-made habitats, on the other hand, are typically more environmentally homogeneous and generally contain plants that are pre-adapted to disturbance and warm and dry conditions (Kalusová et al., 2017). Therefore, it is expected that human-made habitats would often contain plant taxa characterized by relatively large *SLA* and small *SM*, irrespective of macroclimate. Nevertheless, this result once again underlines the importance of incorporating habitat information into macroecological analyses, as failing to do so might blur the environmental signals that exist at a particular spatial scale.

### 4.2 Projected change in maximum height

Our results showed that the hierarchy of environmental drivers shaping the large-scale distribution of plant height is largely habitat-specific and that the trait patterns found across non-native plant assemblages often deviate from those found in native assemblages. Specifically, temperature plays the dominant role in the projected change of *H*_max_ in forests and heaths and scrub, while in other habitats, the effect of precipitation prevails. Similarly, a strong positive association between temperature and plant height has been detected across habitat types and within specific habitats at the continental (e.g. Šímová et al., 2018; forest understories, Padullés Cubino et al., 2021) and regional scales (e.g., forest understories, Maes et al., 2020; mountain grasslands and rock and scree, Dubuis et al., 2013). Moles et al. (2009), on the other hand, reported that the best predictor of plant height at the global scale is the precipitation of the wettest month, which is supported by our results for *H*_max_ in grasslands and rock and scree habitats. Interestingly, while the relationships for native woody assemblages add to the current consensus on the effect of climate on woody plant height (Šímová et al., 2018; Swenson et al., 2012), the results for neophytes may suggest that woody non-natives are preadapted to a different subgradient of the global climatic gradient – in particular, to hotter and drier conditions – where the opposite, negative relationship with temperature can occur (Madani et al., 2018; Moles, 2018). A negative relationship between *H*_max_ of invasive neophytes and *T*_cold_ was also shown by Milanović et al. (2020), although the mechanisms behind this phenomenon remain unclear.

### 4.3 Projected change in specific leaf area

Contrary to several previous studies (Dubuis et al., 2013; Rosbakh et al., 2015; Šímová et al., 2018), herbaceous native and neophyte *SLA* correlated primarily negatively with temperature, and only herbaceous archaeophyte *SLA* tended to show the opposite. Somewhat unexpectedly, another pronounced driver of herbaceous *SLA* change, *P*_dry_, also had mainly a negative effect on *SLA* across all herbaceous assemblages, whereas *P*_wet_ had a positive and negative influence on herbaceous native and neophyte *SLA*, respectively (Figure 5). A negative shift of native *SLA* along the *P*_dry_ gradient appears in disagreement with a previously documented negative effect of drought on *SLA* (Wellstein et al., 2017; Wright et al., 2005). This pattern may be attributed to the fact that within Central Europe, plants with evergreen, low-SLA leaves predominantly occur in the mountains (where precipitation is high) (Chytrý et al., 2021). In contrast to herbaceous *SLA* yet in alignment with previous reports (Šímová et al., 2018; Swenson et al., 2012), *SLA* across all woody assemblages exhibited a strong positive relationship with temperature as well as *P*_dry_. As a result, it is predicted that environmental change will lead to an increase in woody native *SLA* and even more so in woody neophyte *SLA* (Figures 3b,d, 5, S5), which may allow non-natives to gain a further advantage over natives (Pyšek & Richardson, 2008).

Reflecting the combined effect of all the predictors, our projections forecasted mostly a decrease in herbaceous *SLA*; an increase throughout the study area was projected only for native herbaceous *SLA* in wetlands and archaeophyte *SLA* in forests, and a partial increase in *SLA* was projected for archaeophytes in wetlands and human-made habitats and for herbaceous neophytes in grasslands and human-made habitats (Figures 3, S4). Despite a general projected shift towards more conservative resource-use strategies, our projections suggest that herbaceous neophyte *SLA* will be affected less than that of herbaceous natives (Figures S4, S16). This may lead to a further increase in the *SLA* imbalance between native and non-native taxa towards the latter in the region (Divíšek et al., 2018), possibly resulting in an even higher proportion of invasive non-natives (Pyšek & Richardson, 2008). The alteration of the *SLA* composition will undoubtedly affect ecosystem functioning. For example, an overall decrease of *SLA* in grasslands may lead to higher root biomass (Klimešová et al., 2021) and total soil carbon (Garnier et al., 2004), as well as reduced nutrient cycling (Lavorel et al., 2011) and productivity (Brun et al., 2022).

### 4.4 Projected change in seed mass

Our results show that overall drier, less stable climates may on average contribute to an increase in herbaceous *SM* but a decrease in woody *SM*. This finding is congruent with previous studies (Baker, 1972; Dubuis et al., 2013; Šímová et al., 2015; Swenson et al., 2012; Vandelook et al., 2018) and at least partially explains the heterogeneous relationships of *SM* and precipitation in the literature (discussed in Moles, 2018). Notably, while this pattern holds for natives and archaeophytes across all habitats, neophytes often deviate from it.

Specifically, we observed the opposite effect of the precipitation amount and seasonality on herbaceous neophyte *SM* in tree- and shrub-dominated as well as rock and scree habitats and on woody neophyte *SM* in human-made habitats (Figures 5, S14). Such divergence from native herbaceous *SM* may be confounded with the turnover in the growth form, i.e., the proportional increase of small-seeded, short-leaved neophytes in drier areas (Sandel et al., 2010). Additionally, the contrasting responses of native and non-native plant assemblages to precipitation may be due to the differences in the duration of their exposure to the environment and the fact that many non-natives are still actively spreading. For archaeophytes, the positive effect of precipitation is likely to be overwhelmed by the negative effect of higher temperatures, leading to the overall decrease in *SM* (Figures 4, S6). Importantly, for woody *SM*, the overall effect of precipitation may be not as pronounced as that of temperature. Particularly, our results point to a strong positive association of woody *SM* with *T*_warm_ and *T*_cold_ (Figure 5), which drives the projected increase in woody *SM* (Figures 4b,d, S7). This is in line with previous studies, which also documented a strong positive effect of temperature on *SM* of woody plants (Moles et al., 2014; Šímová et al., 2015, 2018; Swenson et al., 2012) and highlighted that herbaceous *SM* is less sensitive to temperature than woody *SM* (Šímová et al., 2018).

### 4.5 Differences in the response between native and non-native taxa

The observed differences between native and non-native plant assemblages once again point out that biogeographical origin affects species performance, via eco-evolutionary novelty of non-natives (Heger et al., 2019; Saul et al., 2013) or due to their pre-adaptation to specific conditions (Maron et al., 2004). For example, non-native species often originate from more nitrogen-rich habitats (Dostál et al., 2013) and therefore are characterized by higher *SLA*. In the long run, the differential response of native and non-native species to environmental factors might lead to even stringer differences in their trait compositions, and this is especially likely for neophytes. Such differences suggest that ecosystem functions provided by future neophyte assemblages may not be redundant to those currently provided by natives. On the contrary, functions currently provided by natives may be replaced with different functions provided by neophytes, thus leading to the increase of functional turnover rather than its buffering in the course of global change.

### 4.6 Study shortcomings

In our analyses, we aggregated plant trait data to the taxon level and then to the 10’ × 6’ grid-cell level for a total of 24 unique assemblages defined by plant woodiness, biogeographical status, and habitat type. Although this precluded us from incorporating intraspecific trait variation, the resolution of trait data was sufficient to accurately model the trait variation due to species sorting as the function of environmental predictors. Moreover, we ensured that trait data were representative of our study area by primarily using data sources for Central Europe and where possible excluding data from environments not found in Germany or the Czech Republic. Additionally, the decisions made during data preparation (e.g., the use of geometric vs. arithmetic mean for trait averaging) might have to a certain extent influenced our results, especially for non-native plants, whose number per grid cell was typically low. This problem, however, was to some extent ameliorated by partial pooling of parameter estimates for different plant assemblages within the multilevel modeling framework. We generated trait change projections using Bayesian stacking of predictive distributions from individual-predictor models (Yao et al., 2018), instead of parametrizing predictive models with all environmental predictors at once. In our case, the latter was difficult to implement because the same categorical variables (i.e., woodiness and biogeographical status) would be included in many interaction terms with intercorrelated environmental predictors (Figure S19), which might have led to spurious results (Duncan & Kefford, 2021). Finally, the obtained projections of future trait change were highly uncertain (Figures S8–13), which indicates that other important predictors of trait composition need to be identified and incorporated into further analyses (e.g., soil variables, Joswig et al., 2022).

### 4.7 Conclusions

In this study, we assessed how habitat-specific plant trait values might shift under future environmental change. Our results depicted the frequently neglected distinctions in trait–environment relationships that are contingent upon plant woodiness, biogeographical status, and habitat type, thereby explaining some of the existing idiosyncrasies within the literature and producing more informative and refined projections of future trait changes. The obtained projections, although uncertain and requiring more global change drivers to factor in, provide an insightful perspective on the conditions under which non-native plants may prevail over natives and vice versa and can serve as a starting point for exploring changes in ecosystem functions and services in a rapidly changing world.

## ACKNOWLEDGEMENTS

We thank Milan Chytrý and Hanno Seebens for their valuable comments on the earlier versions of the manuscript. This research was part of the AlienScenarios project, funded through the 2017–2018 Belmont Forum and BiodivERsA joint call for research proposals, under the BiodivScen ERA-Net COFUND program, and with the funding organizations BMBF (project no. 01LC1807C – MG, IK, SK), the Austrian Science Foundation (FWF project no. I 4011-B32 – FE, BeL, GL) and the Canadian Natural Sciences and Engineering Research Council grant (BrL). The scientific results were partially computed at the High-Performance Computing (HPC) Cluster EVE, a joint effort of the Helmholtz Centre for Environmental Research – UFZ (www.ufz.de) and the German Centre for Integrative Biodiversity Research (iDiv) Halle-Jena-Leipzig (www.idiv-biodiversity.de).

## AUTHOR CONTRIBUTIONS

IK and SK conceived the initial idea of the study, which was further developed by MG. MG collected the data, performed statistical analyses, and wrote the first draft of the manuscript. All authors contributed critically to the drafts and gave final approval for publication.

## DATA AVAILABILITY STATEMENT

All data used in this study are freely available online.

## ORCID

Marina Golivets https://orcid.org/0000-0003-1278-2663

Ingolf Kühn https://orcid.org/0000-0003-1691-8249

Sonja Knapp https://orcid.org/0000-0001-5792-4691

Bernd Lenzner https://orcid.org/0000-0002-2616-3479

Guillaume Latombe https://orcid.org/0000-0002-8589-8387

Franz Essl https://orcid.org/0000-0001-8253-2112

Brian Leung https://orcid.org/0000-0002-8323-9628

