## Supporting Information for "Projecting the futures of plant traits across habitats in Central Europe"

**Appendix S1:** Original references for the data in the TRY database

Ackerly, D. D., & Cornwell, W. K. (2007). A trait‐based approach to community assembly: partitioning of species trait values into within‐and among‐community components. Ecology Letters, 10(2), 135-145.

Adler, P. B., Salguero-Gómez, R., Compagnoni, A., Hsu, J. S., Ray-Mukherjee, J., Mbeau-Ache, C., & Franco, M. (2014). Functional traits explain variation in plant life history strategies. Proceedings of the National Academy of Sciences, 111(2), 740-745.

Atkin, O. K., Westbeek, M. H., Cambridge, M. L., Lambers, H., & Pons, T. L. (1997). Leaf respiration in light and darkness (a comparison of slow-and fast-growing *Poa* species). Plant Physiology, 113(3), 961-965.

Atkin, O. K., Schortemeyer, M., McFarlane, N., & Evans, J. R. (1999). The response of fast-and slow-growing Acacia species to elevated atmospheric CO2: an analysis of the underlying components of relative growth rate. Oecologia, 120(4), 544-554.

Baruch, Z., & Goldstein, G. (1999). Leaf construction cost, nutrient concentration, and net CO2 assimilation of native and invasive species in Hawaii. Oecologia, 121(2), 183-192.

Belluau, M., & Shipley, B. (2018). Linking hard and soft traits: Physiology, morphology and anatomy interact to determine habitat affinities to soil water availability in herbaceous dicots. Plos one, 13(3), e0193130.

Blonder, B., Buzzard, B., Sloat, L., Simova, I., Lipson, R., Boyle, B., & Enquist, B. (2012). The shrinkage effect biases estimates of paleoclimate. American Journal of Botany, 99, 1756-1763.

Blonder, B., Violle, C., & Enquist, B. J. (2013). Assessing the causes and scales of the leaf economics spectrum using venation networks in *Populus tremuloides*. Journal of Ecology, 101(4), 981-989.

Blonder, B., Violle, C., Bentley, L. P., & Enquist, B. J. (2011). Venation networks and the origin of the leaf economics spectrum. Ecology letters, 14(2), 91-100.

Bond-Lamberty, B., Wang, C., & Gower, S. T. (2002). Aboveground and belowground biomass and sapwood area allometric equations for six boreal tree species of northern Manitoba. Canadian Journal of Forest Research, 32(8), 1441-1450.

Bond-Lamberty, B., Wang, C., Gower, S. T., & Norman, J. (2002). Leaf area dynamics of a boreal black spruce fire chronosequence. Tree physiology, 22(14), 993-1001.

Bond-Lamberty, B., Wang, C., & Gower, S. T. (2003). The use of multiple measurement techniques to refine estimates of conifer needle geometry. Canadian Journal of Forest Research, 33(1), 101-105.

Bond‐Lamberty, B., Wang, C., & Gower, S. T. (2004). Net primary production and net ecosystem production of a boreal black spruce wildfire chronosequence. Global Change Biology, 10(4), 473-487.

Bragazza, L. (2009). Conservation priority of Italian Alpine habitats: a floristic approach based on potential distribution of vascular plant species. Biodiversity and conservation, 18(11), 2823-2835.

Burrascano, S., Copiz, R., Del Vico, E., Fagiani, S., Giarrizzo, E., Mei, M., ... & Blasi, C. (2015). Wild boar rooting intensity determines shifts in understorey composition and functional traits. Community Ecology, 16(2), 244-253.

Cadotte, M. W. (2017). Functional traits explain ecosystem function through opposing mechanisms. Ecology Letters, 20(8), 989-996.

Campbell, C., Atkinson, L., Zaragoza‐Castells, J., Lundmark, M., Atkin, O., & Hurry, V. (2007). Acclimation of photosynthesis and respiration is asynchronous in response to changes in temperature regardless of plant functional group. New Phytologist, 176(2), 375-389.

Campetella, G., Botta-Dukát, Z., Wellstein, C., Canullo, R., Gatto, S., Chelli, S., ... & Bartha, S. (2011). Patterns of plant trait–environment relationships along a forest succession chronosequence. Agriculture, ecosystems & environment, 145(1), 38-48.

Carswell, F. E., Meir, P., Wandelli, E. V., Bonates, L. C. M., Kruijt, B., Barbosa, E. M., ... & Jarvis, P. G. (2000). Photosynthetic capacity in a central Amazonian rain forest. Tree physiology, 20(3), 179-186.

Cavender-Bares, J., Keen, A., & Miles, B. (2006). Phylogenetic structure of Floridian plant communities depends on taxonomic and spatial scale. Ecology, 87(sp7), S109-S122.

Choat, B., Jansen, S., Brodribb, T. J., Cochard, H., Delzon, S., Bhaskar, R., ... & Zanne, A. E. (2012). Global convergence in the vulnerability of forests to drought. Nature, 491(7426), 752-755.

Ciccarelli, D. (2015). Mediterranean coastal dune vegetation: are disturbance and stress the key selective forces that drive the psammophilous succession?. Estuarine, Coastal and Shelf Science, 165, 247-253.

Ciocarlan V. (2009). The illustrated Flora of Romania. Pteridophyta et Spermatopyta. Editura Ceres, 1141 p.

Cornelissen, J. H. C. (1996). An experimental comparison of leaf decomposition rates in a wide range of temperate plant species and types. Journal of ecology, 573-582.

Cornelissen, J. H. C., Cerabolini, B., Castro‐Díez, P., Villar‐Salvador, P., Montserrat‐Martí, G., Puyravaud, J. P., ... & Aerts, R. (2003). Functional traits of woody plants: correspondence of species rankings between field adults and laboratory‐grown seedlings?. Journal of Vegetation Science, 14(3), 311-322.

Cornelissen, J. H. C., Diez, P. C., & Hunt, R. (1996). Seedling growth, allocation and leaf attributes in a wide range of woody plant species and types. Journal of ecology, 755-765.

Cornelissen, J. H., Pérez-Harguindeguy, N., Díaz, S., Grime, J. P., Marzano, B., Cabido, M., ... & Cerabolini, B. (1999). Leaf structure and defence control litter decomposition rate across species and life forms in regional floras on two continents. The New Phytologist, 143(1), 191-200.

Cornelissen, J., Aerts, R., Cerabolini, B., Werger, M., & Van Der Heijden, M. (2001). Carbon cycling traits of plant species are linked with mycorrhizal strategy. Oecologia, 129(4), 611-619.

Cornwell, W. K., & Ackerly, D. D. (2009). Community assembly and shifts in plant trait distributions across an environmental gradient in coastal California. Ecological Monographs, 79(1), 109-126.

Cornwell, W. K., Schwilk, D. W., & Ackerly, D. D. (2006). A trait‐based test for habitat filtering: convex hull volume. Ecology, 87(6), 1465-1471.

Craine, J. M., Nippert, J. B., Towne, E., Tucker, S., Kembel, S. W., Skibbe, A., & McLauchlan, K. K. (2011). Functional consequences of climate change-induced plant species loss in a tallgrass prairie. Oecologia, 165(4), 1109-1117.

Craine, J. M., Ocheltree, T. W., Nippert, J. B., Towne, E., Skibbe, A. M., Kembel, S. W., & Fargione, J. E. (2013). Global diversity of drought tolerance and grassland climate-change resilience. Nature Climate Change, 3(1), 63-67.

Craine, J. M., Towne, E., Ocheltree, T. W., & Nippert, J. B. (2012). Community traitscape of foliar nitrogen isotopes reveals N availability patterns in a tallgrass prairie. Plant and Soil, 356(1), 395-403.

Dainese, M., & Bragazza, L. (2012). Plant traits across different habitats of the Italian Alps: a comparative analysis between native and alien species. Alpine Botany, 122(1), 11-21.

Dalke, I. V., Novakovskiy, A. B., Maslova, S. P., & Dubrovskiy, Y. A. (2018). Morphological and functional traits of herbaceous plants with different functional types in the European Northeast. Plant Ecology, 219(11), 1295-1305.

de Frutos, A., Navarro, T., Pueyo, Y., & Alados, C. L. (2015). Inferring resilience to fragmentation-induced changes in plant communities in a semi-arid Mediterranean ecosystem. PloS one, 10(3), e0118837.

de Vries, F. T., & Bardgett, R. D. (2016). Plant community controls on short‐term ecosystem nitrogen retention. New Phytologist, 210(3), 861-874.

Diaz, S., Hodgson, J. G., Thompson, K., Cabido, M., Cornelissen, J. H., Jalili, A., ... & Zak, M. R. (2004). The plant traits that drive ecosystems: evidence from three continents. Journal of vegetation science, 15(3), 295-304.

Dwyer, J. M., Hobbs, R. J., & Mayfield, M. M. (2014). Specific leaf area responses to environmental gradients through space and time. Ecology, 95(2), 399-410.

Fagúndez, J., Juan, R., Fernández, I., Pastor, J., & Izco, J. (2010). Systematic relevance of seed coat anatomy in the European heathers (Ericeae, Ericaceae). Plant Systematics and Evolution, 284(1), 65-76.

Falster, D. S., Duursma, R. A., Ishihara, M. I., Barneche, D. R., FitzJohn, R. G., Vårhammar, A., ... & York, R. A. (2015). BAAD: A biomass and allometry database for woody plants. Ecology 96:1445.

Fonseca, C. R., Overton, J. M., Collins, B., & Westoby, M. (2000). Shifts in trait‐combinations along rainfall and phosphorus gradients. Journal of Ecology, 88(6), 964-977.

Freschet, G. T., Cornelissen, J. H., van Logtestijn, R. S., & Aerts, R. (2010). Substantial nutrient resorption from leaves, stems and roots in a subarctic flora: what is the link with other resource economics traits?. New Phytologist, 186(4), 879-889.

Gachet, S., Véla, E., & Tatoni, T. (2005). BASECO: a floristic and ecological database of Mediterranean French flora. Biodiversity & Conservation, 14(4), 1023-1034.

Giarrizzo, E., Burrascano, S., Chiti, T., de Bello, F., Lepš, J., Zavattero, L., & Blasi, C. (2017). Re‐visiting historical semi‐natural grasslands in the Apennines to assess patterns of changes in species composition and functional traits. Applied Vegetation Science, 20(2), 247-258.

Gonzalez-Akre, E., McShea, W., Bourg, N., Anderson-Teixeira, K. 2015. Leaf traits data (SLA) for 56 woody species at the Smithsonian Conservation Biology Institute-ForestGEO Forest Dynamic Plot. Front Royal, Virginia. USA. [Data set]. Version 1.0.

Gos, P., Loucougaray, G., Colace, M. P., Arnoldi, C., Gaucherand, S., Dumazel, D., ... & Lavorel, S. (2016). Relative contribution of soil, management and traits to co-variations of multiple ecosystem properties in grasslands. Oecologia, 180(4), 1001-1013.

Green, W. 2009. USDA PLANTS Compilation, version 1, 09-02-02. Available from: http://bricol.net/downloads/data/PLANTSdatabase/

Guy, A. L., Mischkolz, J. M., & Lamb, E. G. (2013). Limited effects of simulated acidic deposition on seedling survivorship and root morphology of endemic plant taxa of the Athabasca Sand Dunes in well-watered greenhouse trials. Botany, 91(3), 176-181.

Herz, K., Dietz, S., Haider, S., Jandt, U., Scheel, D., & Bruelheide, H. (2017). Drivers of intraspecific trait variation of grass and forb species in German meadows and pastures. Journal of Vegetation Science, 28(4), 705-716.

Herz, K., Dietz, S., Haider, S., Jandt, U., Scheel, D., & Bruelheide, H. (2017). Predicting individual plant performance in grasslands. Ecology and evolution, 7(21), 8958-8965.

Kattge, J., Knorr, W., Raddatz, T., & Wirth, C. (2009). Quantifying photosynthetic capacity and its relationship to leaf nitrogen content for global‐scale terrestrial biosphere models. Global Change Biology, 15(4), 976-991.

Kazakou, E., Vile, D., Shipley, B., Gallet, C., & Garnier, E. (2006). Co‐variations in litter decomposition, leaf traits and plant growth in species from a Mediterranean old‐field succession. Functional Ecology, 20(1), 21-30.

Kichenin, E., Wardle, D. A., Peltzer, D. A., Morse, C. W., & Freschet, G. T. (2013). Contrasting effects of plant inter‐and intraspecific variation on community‐level trait measures along an environmental gradient. Functional Ecology, 27(5), 1254-1261.

La Pierre, K. J., & Smith, M. D. (2015). Functional trait expression of grassland species shift with short-and long-term nutrient additions. Plant Ecology, 216(2), 307-318.

Laughlin, D. C., Leppert, J. J., Moore, M. M., & Sieg, C. H. (2010). A multi‐trait test of the leaf‐height‐seed plant strategy scheme with 133 species from a pine forest flora. Functional Ecology, 24(3), 493-501.

Laughlin, D. C., Fule, P. Z., Huffman, D. W., Crouse, J., & Laliberté, E. (2011). Climatic constraints on trait‐based forest assembly. Journal of Ecology, 99(6), 1489-1499.

Lavorel, S., Grigulis, K., Lamarque, P., Colace, M. P., Garden, D., Girel, J., ... & Douzet, R. (2011). Using plant functional traits to understand the landscape distribution of multiple ecosystem services. Journal of Ecology, 99(1), 135-147.

Letts, B., Lamb, E. G., Mischkolz, J. M., & Romo, J. T. (2015). Litter accumulation drives grassland plant community composition and functional diversity via leaf traits. Plant Ecology, 216(3), 357-370.

Lhotsky, B., Csecserits, A., Kovács, B., & Botta-Dukát, Z. (2016). New plant trait records of the Hungarian flora. Acta Botanica Hungarica, 58(3-4), 397-400.

Li, Y., & Shipley, B. (2018). Community divergence and convergence along experimental gradients of stress and disturbance. Ecology, 99, 775-781.

Liebergesell, M., Reu, B., Stahl, U., Freiberg, M., Welk, E., Kattge, J., ... & Wirth, C. (2016). Functional resilience against climate-driven extinctions–Comparing the functional diversity of European and North American tree floras. PloS one, 11(2), e0148607.

Louault, F., Pillar, V. D., Aufrere, J., Garnier, E., & Soussana, J. F. (2005). Plant traits and functional types in response to reduced disturbance in a semi‐natural grassland. Journal of vegetation Science, 16(2), 151-160.

Loveys, B. R., Atkinson, L. J., Sherlock, D. J., Roberts, R. L., Fitter, A. H., & Atkin, O. K. (2003). Thermal acclimation of leaf and root respiration: an investigation comparing inherently fast‐and slow‐growing plant species. Global Change Biology, 9(6), 895-910.

Lukeš, P., Stenberg, P., Rautiainen, M., Mõttus, M., & Vanhatalo, K. M. (2013). Optical properties of leaves and needles for boreal tree species in Europe. Remote Sensing Letters, 4(7), 667-676.

Maire, V., Wright, I. J., Prentice, I. C., Batjes, N. H., Bhaskar, R., Van Bodegom, P. M., ... & Santiago, L. S. (2015). Global effects of soil and climate on leaf photosynthetic traits and rates. Global Ecology and Biogeography, 24(6), 706-717.

Manning, P., Newington, J. E., Robson, H. R., Saunders, M., Eggers, T., Bradford, M. A., ... & Rees, M. (2006). Decoupling the direct and indirect effects of nitrogen deposition on ecosystem function. Ecology Letters, 9(9), 1015-1024.

Martin, A. R., Hale, C. E., Cerabolini, B. E., Cornelissen, J. H., Craine, J., Gough, W. A., ... & Tirona, C. K. (2018). Inter-and intraspecific variation in leaf economic traits in wheat and maize. AoB Plants, 10(1), ply006.

McDonald, P. G., Fonseca, C. R., McC, J., & Westoby, M. (2003). Leaf-size divergence along rainfall and soil-nutrient gradients: is the method of size reduction common among clades?. Functional Ecology, 50-57.

McKenna, M. F., & Shipley, B. (1999). Interacting determinants of interspecific relative growth: empirical patterns and a theoretical explanation. Ecoscience, 6(2), 286-296.

McPartland M. Alaska Peatland Experiment (APEX) 2016 PFT values.

Medeiros, J. S., Burns, J. H., Nicholson, J., Rogers, L., & Valverde‐Barrantes, O. (2017). Decoupled leaf and root carbon economics is a key component in the ecological diversity and evolutionary divergence of deciduous and evergreen lineages of genus Rhododendron. American Journal of Botany, 104(6), 803-816.

Medlyn, B. E., & Jarvis, P. G. (1999). Design and use of a database of model parameters from elevated [CO2] experiments. Ecological Modelling, 124(1), 69-83.

Medlyn, B. E., Barton, C. V. M., Broadmeadow, M. S. J., Ceulemans, R., De Angelis, P., Forstreuter, M., ... & Jarvis, P. G. (2001). Stomatal conductance of forest species after long‐term exposure to elevated CO2 concentration: a synthesis. New Phytologist, 149(2), 247-264.

Medlyn, B. E., Badeck, F. W., De Pury, D. G. G., Barton, C. V. M., Broadmeadow, M., Ceulemans, R., ... & Jstbid, P. G. (1999). Effects of elevated [CO2] on photosynthesis in European forest species: a meta‐analysis of model parameters. Plant, Cell & Environment, 22(12), 1475-1495.

Meir, P., Levy, P. E., Grace, J., & Jarvis, P. G. (2007). Photosynthetic parameters from two contrasting woody vegetation types in West Africa. Plant Ecology, 192(2), 277-287.

Meir, P., Kruijt, B., Broadmeadow, M., Barbosa, E., Kull, O., Carswell, F., ... & Jarvis, P. G. (2002). Acclimation of photosynthetic capacity to irradiance in tree canopies in relation to leaf nitrogen concentration and leaf mass per unit area. Plant, cell & environment, 25(3), 343-357.

Meng, T. T., Wang, H., Harrison, S. P., Prentice, I. C., Ni, J., & Wang, G. (2015). Responses of leaf traits to climatic gradients: adaptive variation versus compositional shifts. Biogeosciences, 12(18), 5339-5352.

Meziane, D., & Shipley, B. (1999). Interacting components of interspecific relative growth rate: constancy and change under differing conditions of light and nutrient supply. Functional Ecology, 13(5), 611-622.

Meziane, D., & Shipley, B. (1999). Interacting determinants of specific leaf area in 22 herbaceous species: effects of irradiance and nutrient availability. Plant, Cell & Environment, 22(5), 447-459.

Beckmann, M., Hock, M., Bruelheide, H., & Erfmeier, A. (2012). The role of UV-B radiation in the invasion of Hieracium pilosella—A comparison of German and New Zealand plants. Environmental and Experimental Botany, 75, 173-180.

Michaletz, S. T., & Johnson, E. A. (2006). A heat transfer model of crown scorch in forest fires. Canadian Journal of Forest Research, 36(11), 2839-2851.

Michaletz, S. T., & Johnson, E. A. (2008). A biophysical process model of tree mortality in surface fires. Canadian Journal of Forest Research, 38(7), 2013-2029.

Milla, R., & Reich, P. B. (2011). Multi-trait interactions, not phylogeny, fine-tune leaf size reduction with increasing altitude. Annals of botany, 107(3), 455-465.

Miller, J. E., Ives, A. R., Harrison, S. P., & Damschen, E. I. (2018). Early‐and late‐flowering guilds respond differently to landscape spatial structure. Journal of Ecology, 106(3), 1033-1045.

Mischkolz, J. M. (2013) Selecting and evaluating native forage mixtures for the mixed grass prairie. Master Thesis, University of Saskatchewan, Saskatoon, SK. http://hdl.handle.net/10388/ETD-2013-04-988

Moles, A. T., Ackerly, D. D., Webb, C. O., Tweddle, J. C., Dickie, J. B., Pitman, A. J., & Westoby, M. (2005). Factors that shape seed mass evolution. Proceedings of the National Academy of Sciences, 102(30), 10540-10544.

Moles, A. T., Falster, D. S., Leishman, M. R., & Westoby, M. (2004). Small‐seeded species produce more seeds per square metre of canopy per year, but not per individual per lifetime. Journal of ecology, 92(3), 384-396.

Moles, A. T., Warton, D. I., Warman, L., Swenson, N. G., Laffan, S. W., Zanne, A. E., ... & Leishman, M. R. (2009). Global patterns in plant height. Journal of ecology, 97(5), 923-932.

Onoda, Y., Wright, I. J., Evans, J. R., Hikosaka, K., Kitajima, K., Niinemets, Ü., ... & Westoby, M. (2017). Physiological and structural tradeoffs underlying the leaf economics spectrum. New Phytologist, 214(4), 1447-1463.

Ordonez, J. C., van Bodegom, P. M., Witte, J. P. M., Bartholomeus, R. P., van Hal, J. R., & Aerts, R. (2010). Plant strategies in relation to resource supply in mesic to wet environments: does theory mirror nature?. The American Naturalist, 175(2), 225-239.

Ordoñez, J. C., Van Bodegom, P. M., Witte, J. P. M., Bartholomeus, R. P., Van Dobben, H. F., & Aerts, R. (2010). Leaf habit and woodiness regulate different leaf economy traits at a given nutrient supply. Ecology, 91(11), 3218-3228.

Pahl, A. T., Kollmann, J., Mayer, A., & Haider, S. (2013). No evidence for local adaptation in an invasive alien plant: field and greenhouse experiments tracing a colonization sequence. Annals of Botany, 112(9), 1921-1930.

Paine, C. T., Amissah, L., Auge, H., Baraloto, C., Baruffol, M., Bourland, N., ... & Hector, A. (2015). Globally, functional traits are weak predictors of juvenile tree growth, and we do not know why. Journal of Ecology, 103(4), 978-989.

Paula, S., & Pausas, J. G. (2008). Burning seeds: germinative response to heat treatments in relation to resprouting ability. Journal of Ecology, 96(3), 543-552.

Paula, S., Arianoutsou, M., Kazanis, D., Tavsanoglu, Ç., Lloret, F., Buhk, C., ... & Pausas, J. G. (2009). Fire‐related traits for plant species of the Mediterranean Basin: Ecological Archives E090‐094. Ecology, 90(5), 1420-1420.

Peco, B., de Pablos, I., Traba, J., & Levassor, C. (2005). The effect of grazing abandonment on species composition and functional traits: the case of dehesa grasslands. Basic and applied Ecology, 6(2), 175-183.

Poschlod, P., Kleyer, M., Jackel, A. K., Dannemann, A., & Tackenberg, O. (2003). BIOPOP—A database of plant traits and internet application for nature conservation. Folia Geobotanica, 38(3), 263-271.

Prentice, I. C., Meng, T., Wang, H., Harrison, S. P., Ni, J., & Wang, G. (2011). Evidence of a universal scaling relationship for leaf CO2 drawdown along an aridity gradient. New Phytologist, 190(1), 169-180.

Preston, K. A., Cornwell, W. K., & DeNoyer, J. L. (2006). Wood density and vessel traits as distinct correlates of ecological strategy in 51 California coast range angiosperms. New Phytologist, 170(4), 807-818.

Price, C. A., & Enquist, B. J. (2007). Scaling mass and morphology in leaves: an extension of the WBE model. Ecology, 88(5), 1132-1141.

Price, C. A., Enquist, B. J., & Savage, V. M. (2007). A general model for allometric covariation in botanical form and function. Proceedings of the National Academy of Sciences, 104(32), 13204-13209.

Pyankov, V. I., Kondratchuk, A. V., & Shipley, B. (1999). Leaf structure and specific leaf mass: the alpine desert plants of the Eastern Pamirs, Tadjikistan. The New Phytologist, 143(1), 131-142.

Reich, P. B., Tjoelker, M. G., Pregitzer, K. S., Wright, I. J., Oleksyn, J., & Machado, J. L. (2008). Scaling of respiration to nitrogen in leaves, stems and roots of higher land plants. Ecology letters, 11(8), 793-801.

Rogers, A., Serbin, S. P., Ely, K. S., Sloan, V. L., & Wullschleger, S. D. (2017). Terrestrial biosphere models underestimate photosynthetic capacity and CO2 assimilation in the Arctic. New Phytologist, 216(4), 1090-1103.

Sanda V., Bita-Nicolae, C. D., & Barabas, N. (2003). The flora of spontane and cultivated cormophytes from Romania. Editura “Ion Borcea”, Bacau, 316 p.

Sandel, B., Corbin, J. D., & Krupa, M. (2011). Using plant functional traits to guide restoration: A case study in California coastal grassland. Ecosphere, 2(2), 1-16.

Schroeder‐Georgi, T., Wirth, C., Nadrowski, K., Meyer, S. T., Mommer, L., & Weigelt, A. (2016). From pots to plots: hierarchical trait‐based prediction of plant performance in a mesic grassland. Journal of Ecology, 104(1), 206-218.

Shipley, B. (2002). Trade‐offs between net assimilation rate and specific leaf area in determining relative growth rate: relationship with daily irradiance. Functional ecology, 16(5), 682-689.

Shipley, B. (1989). The use of above-ground maximum relative growth rate as an accurate predictor of whole-plant maximum relative growth rate. Functional Ecology, 771-775.

Shipley, B. (1995). Structured interspecific determinants of specific leaf area in 34 species of herbaceous angiosperms. Functional ecology, 312-319.

Shipley, B., & Lechowicz, M. J. (2000). The functional co-ordination of leaf morphology, nitrogen concentration, and gas exchange in 40 wetland species. Ecoscience, 7(2), 183-194.

Shipley, B., & Parent, M. (1991). Germination responses of 64 wetland species in relation to seed size, minimum time to reproduction and seedling relative growth rate. Functional ecology, 111-118.

Shipley, B., & Vu, T. T. (2002). Dry matter content as a measure of dry matter concentration in plants and their parts. New Phytologist, 153(2), 359-364.

Siefert, A., Fridley, J. D., & Ritchie, M. E. (2014). Community functional responses to soil and climate at multiple spatial scales: when does intraspecific variation matter?. PLoS one, 9(10), e111189.

Smith, N. G., & Dukes, J. S. (2017). LCE: Leaf carbon exchange data set for tropical, temperate, and boreal species of North and Central America. Ecology, 98, 2978.

Smith, S. W., Woodin, S. J., Pakeman, R. J., Johnson, D., & Van Der Wal, R. (2014). Root traits predict decomposition across a landscape‐scale grazing experiment. New Phytologist, 203(3), 851-862.

Sodhi, D. S., Livingstone, S. W., Carboni, M., & Cadotte, M. W. (2019). Plant invasion alters trait composition and diversity across habitats. Ecology and Evolution, 9(11), 6199-6210.

Spasojevic, M. J., & Suding, K. N. (2012). Inferring community assembly mechanisms from functional diversity patterns: the importance of multiple assembly processes. Journal of Ecology, 100(3), 652-661.

Spasojevic, M. J., Turner, B. L., & Myers, J. A. (2016). When does intraspecific trait variation contribute to functional beta‐diversity?. Journal of Ecology, 104(2), 487-496.

Takkis, K. (2014). Changes in plant species richness and population performance in response to habitat loss and fragmentation. DISSERTATIONES BIOLOGICAE UNIVERSITATIS TARTUENSIS 255, 2014-04-07. Available from: http://hdl.handle.net/10062/39546

Thuiller, W. Traits of European Alpine Flora. OriginAlps Project. Centre National de la Recherche Scientifique.

Tribouillois, H., Fort, F., Cruz, P., Charles, R., Flores, O., Garnier, E., & Justes, E. (2015). A functional characterisation of a wide range of cover crop species: Growth and nitrogen acquisition rates, leaf traits and ecological strategies. PLoS One, 10(3), e0122156.

Tucker, S. S., Craine, J. M., & Nippert, J. B. (2011). Physiological drought tolerance and the structuring of tallgrass prairie assemblages. Ecosphere, 2(4), 1-19.

Vergutz, L., Manzoni, S., Porporato, A., Novais, R. F., & Jackson, R. B. (2012). Global resorption efficiencies and concentrations of carbon and nutrients in leaves of terrestrial plants. Ecological Monographs, 82(2), 205-220.

Vergutz, L., Manzoni, S., Porporato, A., Novais, R. F., & Jackson, R. B. (2012). A global database of carbon and nutrient concentrations of green and senesced leaves. ORNL DAAC. Available from: http://daac.ornl.gov.

Vile, D. (2005). Significations fonctionnelle et ecologique des traits des especes vegetales: exemple dans une succession post-cultural mediterraneenne et generalisations. PhD Thesis.

Walker, A. P., Beckerman, A. P., Gu, L., Kattge, J., Cernusak, L. A., Domingues, T. F., ... & Woodward, F. I. (2014). The relationship of leaf photosynthetic traits–Vcmax and Jmax–to leaf nitrogen, leaf phosphorus, and specific leaf area: a meta‐analysis and modeling study. Ecology and evolution, 4(16), 3218-3235.

Walker, A. P. (2014). A Global Data Set of Leaf Photosynthetic Rates, Leaf N and P, and Specific Leaf Area. ORNL DAAC. Available from: http://daac.ornl.gov.

Wang, H., Harrison, S. P., Prentice, I. C., Yang, Y., Bai, F., Togashi, H. F., ... & Ni, J. (2018). The China plant trait database: Toward a comprehensive regional compilation of functional traits for land plants. Ecology, 99(2), 500.

Wilson, K. B., Baldocchi, D. D., & Hanson, P. J. (2000). Spatial and seasonal variability of photosynthetic parameters and their relationship to leaf nitrogen in a deciduous forest. Tree physiology, 20(9), 565-578.

Wirth, C., & Lichstein, J. W. (2009). The imprint of species turnover on old-growth forest carbon balances-Insights from a trait-based model of forest dynamics. In Old-growth forests (pp. 81-113). Springer, Berlin, Heidelberg.

Wright, J. P., & Sutton‐Grier, A. (2012). Does the leaf economic spectrum hold within local species pools across varying environmental conditions?. Functional Ecology, 26(6), 1390-1398.

Wright, I. J., Reich, P. B., Westoby, M., Ackerly, D. D., Baruch, Z., Bongers, F., ... & Villar, R. (2004). The worldwide leaf economics spectrum. Nature, 428(6985), 821-827.

Wright, I. J., Reich, P. B., Atkin, O. K., Lusk, C. H., Tjoelker, M. G., & Westoby, M. (2006). Irradiance, temperature and rainfall influence leaf dark respiration in woody plants: evidence from comparisons across 20 sites. New Phytologist, 169(2), 309-319.

**TABLE S2**

The combined climate (Representative Concentration Pathways, RCP) and socio-economic scenarios (European Socio-Economic Pathways, Eur-SSP) used in this study. For each RCP, the IMPRESSIONS IAP2 was run with three different dynamically downscaled CMIP5 climate models (i.e., combinations of global climate models, GCM and regional climate models, RCM).

| **RCP** | **GCM** | **RCM** | **Eur-SSP** |
| --- | --- | --- | --- |
| 2.6 | ICHEC-EC-EARTH-r12i1p1 | SMHI-RCA4_v1 | 1, 4 |
| 2.6 | MPI-M-MPI-ESM-LR_r1i1p1 | MPI-CSC-REMO2009_v1 | 1, 4 |
| 2.6 | NCC-NorESM1-M_r1i1p1 | SMHI-RCA4_v1 | 1, 4 |
| 4.5 | NOAA-GFDL-GFDL_ESM2M_r1i1p1 | SMHI_RCA4_v1 | 1, 3, 4 |
| 4.5 | MOHC-HadGEM2-ES_r1i1p1 | SMHI-RCA4_v1 | 1, 3, 4 |
| 4.5 | MPI-M-MPI-ESM-LR_r1i1p1 | CLMcom-CCLM4-8-17_v1 | 1, 3, 4 |
| 8.5 | NOAA-GFDL-GFDL_ESM2M_r1i1p1 | SMHI_RCA4_v1 | 3, 5 |
| 8.5 | MOHC-HadGEM2-ES_r1i1p1 | SMHI-RCA4_v1 | 3, 5 |
| 8.5 | IPSL-IPSL-CM5A-MR_r1i1p1 | IPSL-INERIS-WRF331F_v1 | 3, 5 |

**TABLE S3**

Model predictive performance, measured as the k-fold cross-validation information criterion (*kfoldIC*) and root mean square error (*RMSE*), and model weights used for Bayesian stacking.

| **Environmental predictor** | ***kfoldIC* (± SE)** | ***RMSE*** | **Model weight** |
| --- | --- | --- | --- |
| **Plant maximum height** | | | |
| Precipitation seasonality | 112586.0 ± 414.9 | 0.905 | 0.080 |
| Precipitation of the wettest month | 112367.6 ± 430.4 | 0.902 | 0.314 |
| Temperature of the wettest quarter | 113260.1 ± 410.8 | 0.912 | 0.002 |
| Temperature of the driest quarter | 113337.4 ± 408.7 | 0.913 | 0 |
| Temperature of the coldest month | 112037.8 ± 417.9 | 0.898 | 0.379 |
| Maximum temperature of the warmest month | 113679.8 ± 406.1 | 0.916 | 0.031 |
| % Urban land | 113528.8 ± 403.3 | 0.915 | 0 |
| Precipitation of the driest month | 112861.0 ± 417.9 | 0.907 | 0.193 |
| None (only the CAR structure) | 114152.3 ± 400.6 | 0.922 | - |
| All (weighted average) |  | 0.872 |  |
| **Specific leaf area** | | | |
| Precipitation seasonality | 114118.9 ± 425.7 | 0.915 | 0.014 |
| Precipitation of the wettest month | 113070.3 ± 426.6 | 0.903 | 0.154 |
| Temperature of the wettest quarter | 112889.0 ± 427.3 | 0.902 | 0 |
| Temperature of the driest quarter | 114237.0 ± 421.0 | 0.917 | 0 |
| Temperature of the coldest month | 113108.1 ± 432.3 | 0.904 | 0.262 |
| Maximum temperature of the warmest month | 112387.1 ± 422.5 | 0.896 | 0.335 |
| % Urban land | 117904.6 ± 443.3 | 0.944 | 0 |
| Precipitation of the driest month | 112402.6 ± 422.9 | 0.897 | 0.233 |
| None (only the CAR structure) | 118546.2 ± 438.9 | 0.952 | - |
| All (weighted average) |  | 0.871 |  |
| **Seed mass** | | | |
| Precipitation seasonality | 116673.1 ± 442.1 | 0.932 | 0.275 |
| Precipitation of the wettest month | 118169.3 ± 445.4 | 0.950 | 0.065 |
| Temperature of the wettest quarter | 117851.5 ± 448.1 | 0.944 | 0 |
| Temperature of the driest quarter | 116955.3 ± 440.0 | 0.935 | 0 |
| Temperature of the coldest month | 116837.7 ± 449.1 | 0.933 | 0.281 |
| Maximum temperature of the warmest month | 117914.6 ± 450.5 | 0.944 | 0.164 |
| % Urban land | 117904.6 ± 443.3 | 0.944 | 0 |
| Precipitation of the driest month | 117203.3 ± 443.1 | 0.937 | 0.215 |
| None (only the CAR structure) | 118546.2 ± 438.9 | 0.952 | - |
| All (weighted average) |  | 0.914 |  |

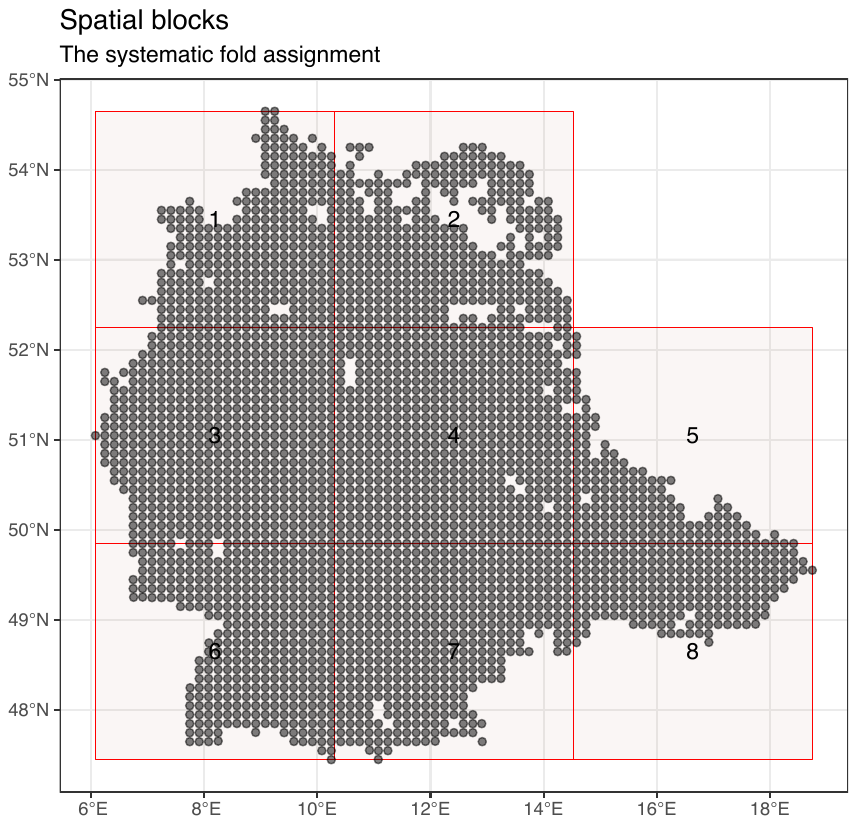

**FIGURE S1**

Spatial block allocation for the k-fold cross-validation. Due to the low sample size, blocks 5 and 8 were merged before analyses, which resulted in 7 blocks total and the number of TK25 grid cells per block ranging from *N* = 301 in block 1 to *N* = 587 in block 4. Grid cells that were used in analyses are shown as grey circles and grid cells that were excluded from analyses due to low sampling effort are shown as gaps.

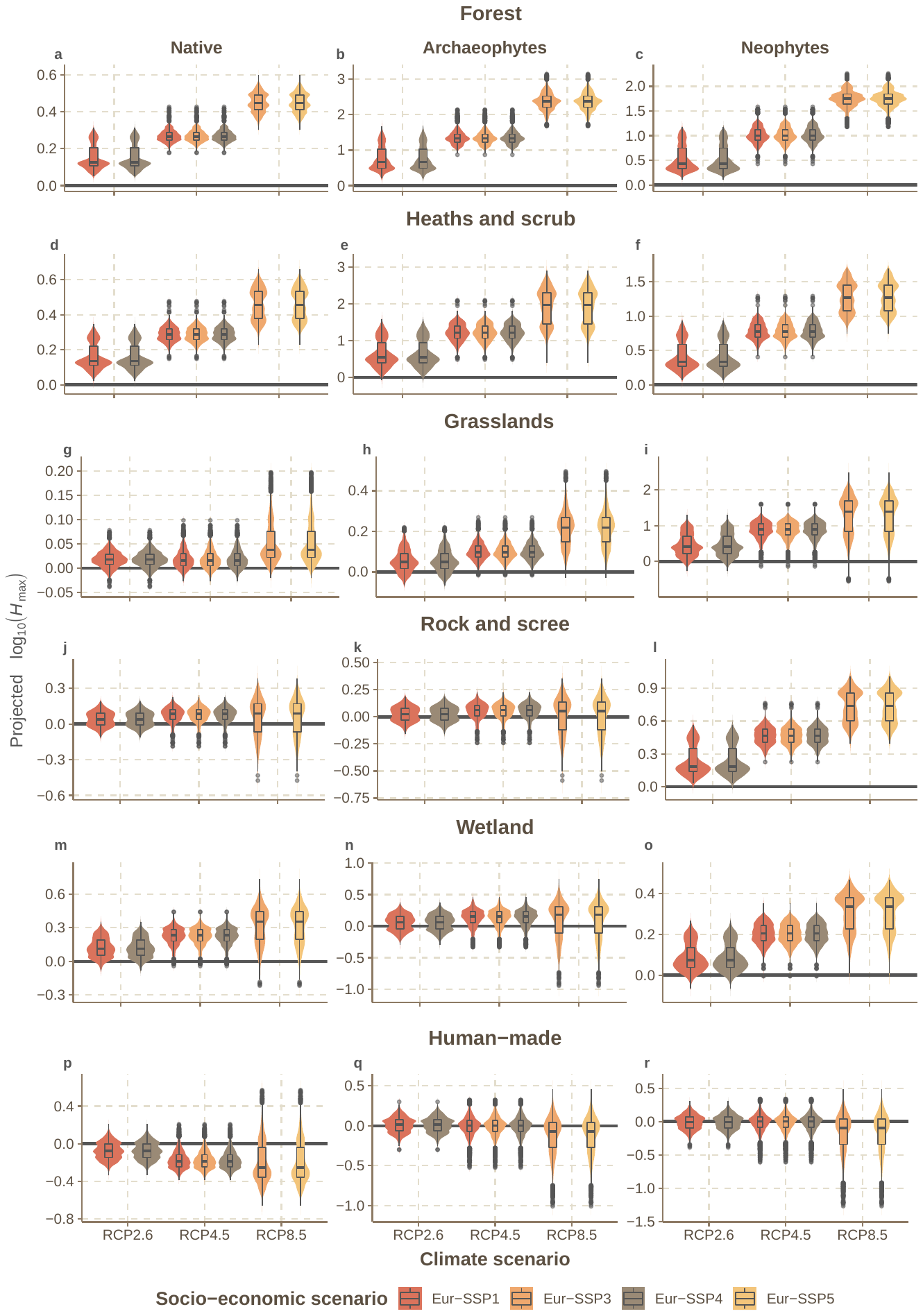

**FIGURE S2**

The projected per-cell (10’ × 10’) change in the log_10_-transformed maximum height (*H*_max_) of *herbaceous* plant assemblages under all plausible combined climate (RCP, all climate models pooled) and socio-economic (Eur-SSP) scenarios for 2081–2100. The trait change here is expressed as the posterior means of per-cell model predictions. The violin plots depict the distributions of projected values across the study area and the boxplots provide summary statistics of those distributions (boxes show 25%, 50%, and 75% quartiles and whiskers give roughly 95% credible intervals). For each habitat by woodiness combination, the trait change is presented in standard deviations (*SD*) of the baseline trait distribution of native species for that combination.

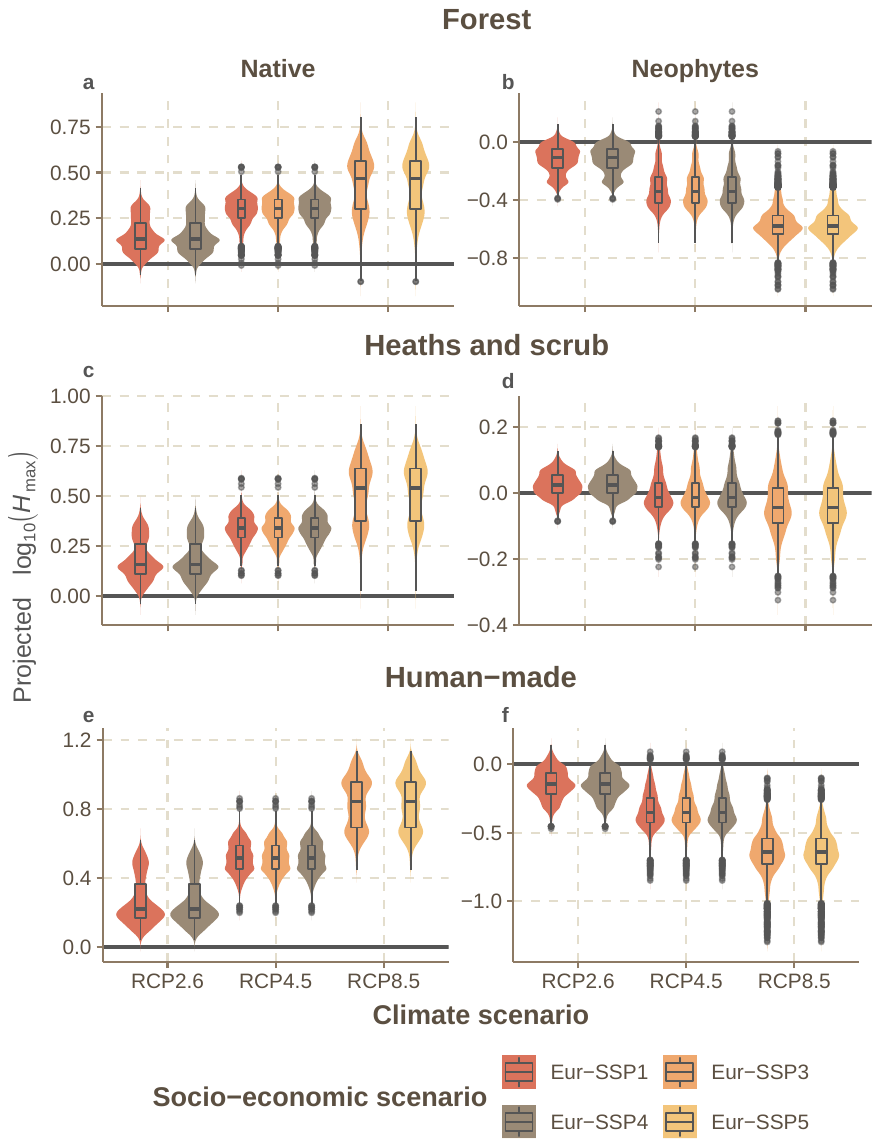

**FIGURE S3**

The projected change in the log_10_-transformed maximum height (*H*_max_) of *woody* plant assemblages under all plausible combined climate (RCP, all climate models pooled) and socio-economic (Eur-SSP) scenarios for 2081–2100. Other details are as in Figure S2.

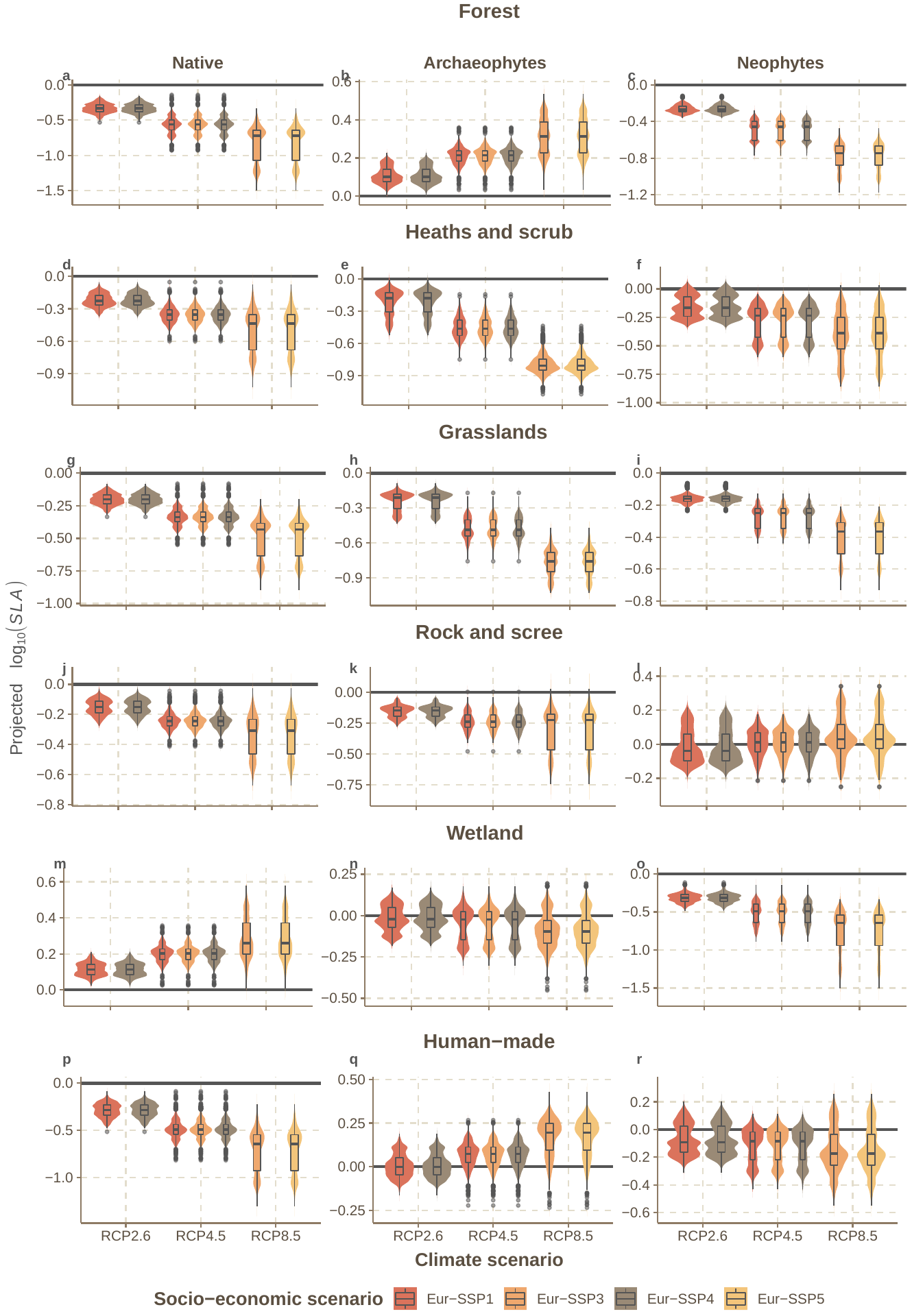

**FIGURE S4**

The projected change in the log_10_-transformed specific leaf area (*SLA*) of *herbaceous* plant assemblages under all plausible combined climate (RCP, all climate models pooled) and socio-economic (Eur-SSP) scenarios for 2081–2100. Other details are as in Figure S2.

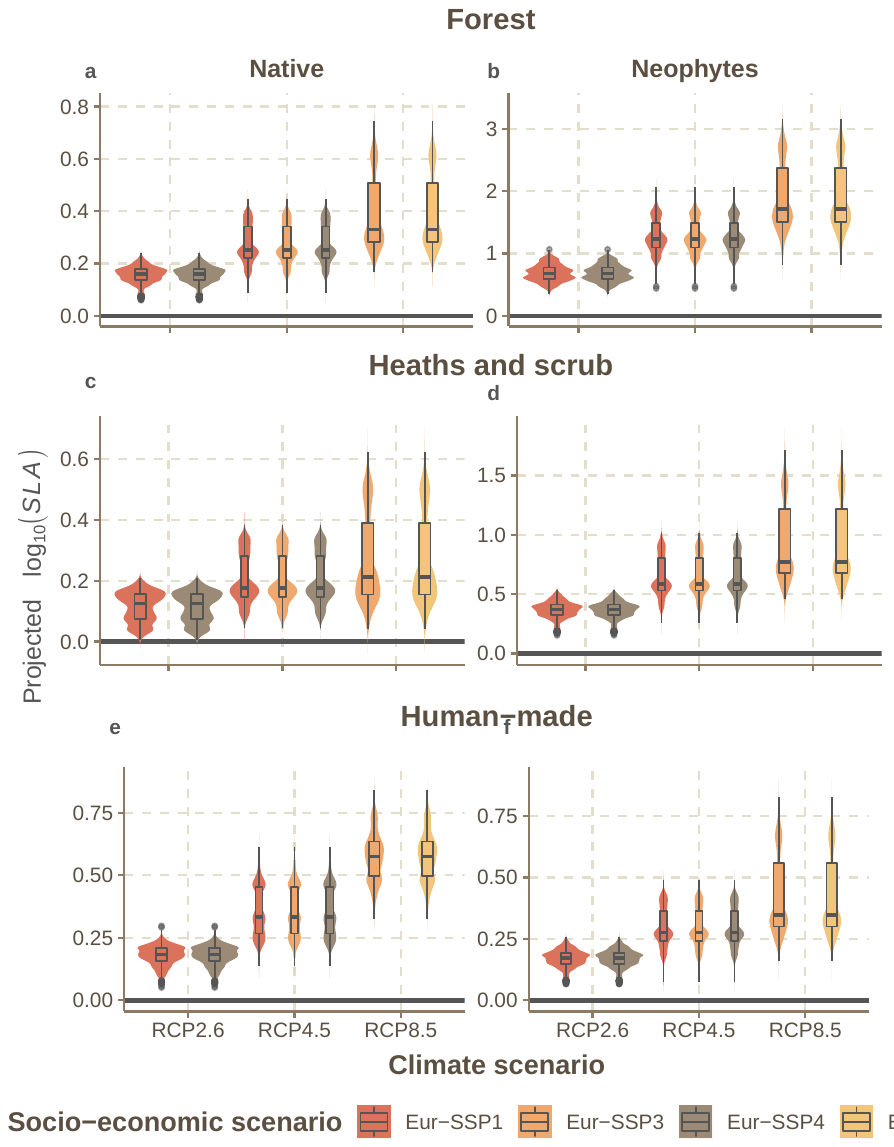

**FIGURE S5**

The projected change in the log_10_-transformed specific leaf area (*SLA*) of *woody* plant assemblages under all plausible combined climate (RCP, all climate models pooled) and socio-economic (Eur-SSP) scenarios for 2081–2100. Other details are as in Figure S2.

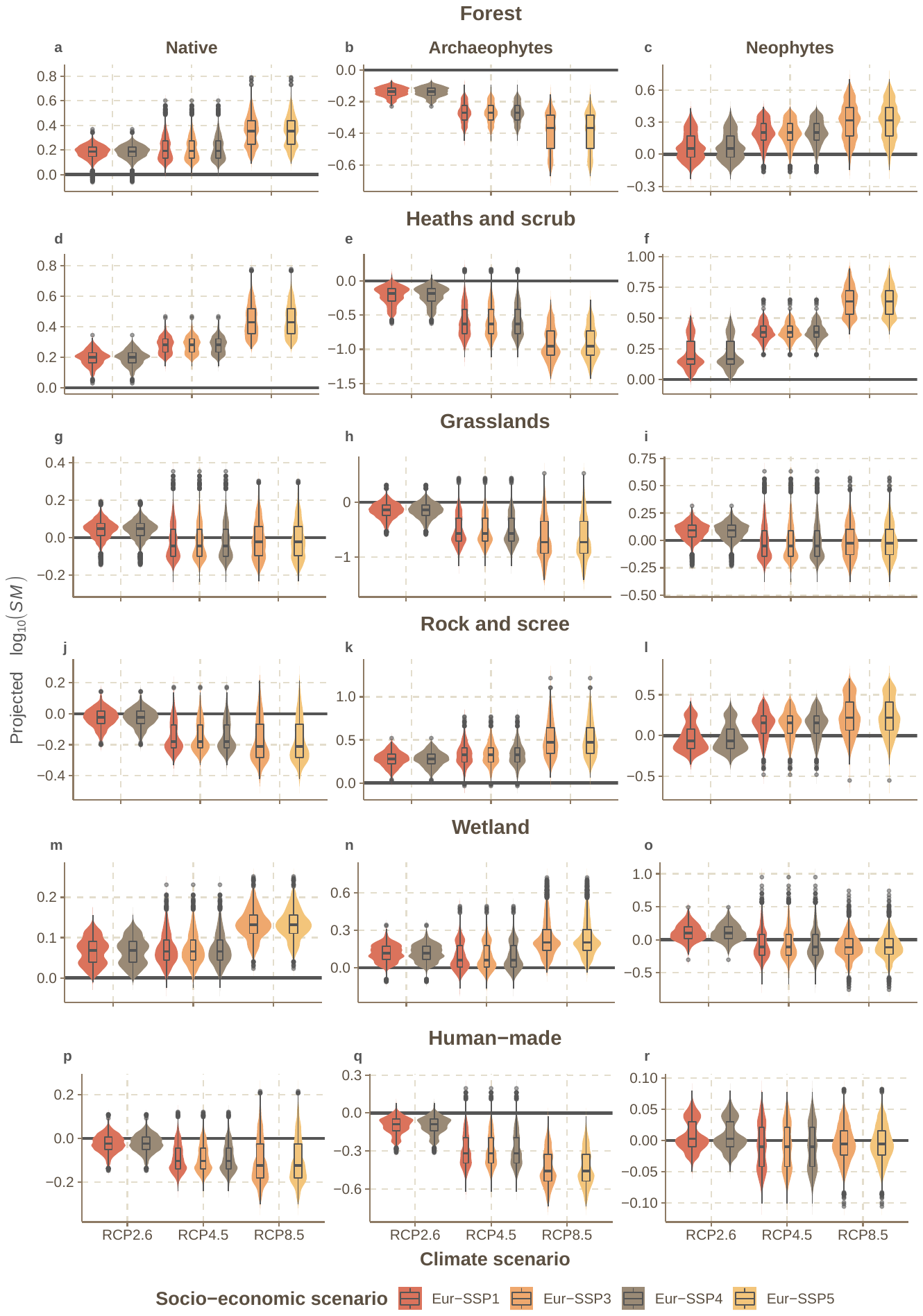

**FIGURE S6**

The projected change in the log_10_-transformed seed mass (*SM*) of *herbaceous* plant assemblages under all plausible combined climate (RCP, all climate models pooled) and socio-economic (Eur-SSP) scenarios for 2081–2100. Other details are as in Figure S2.

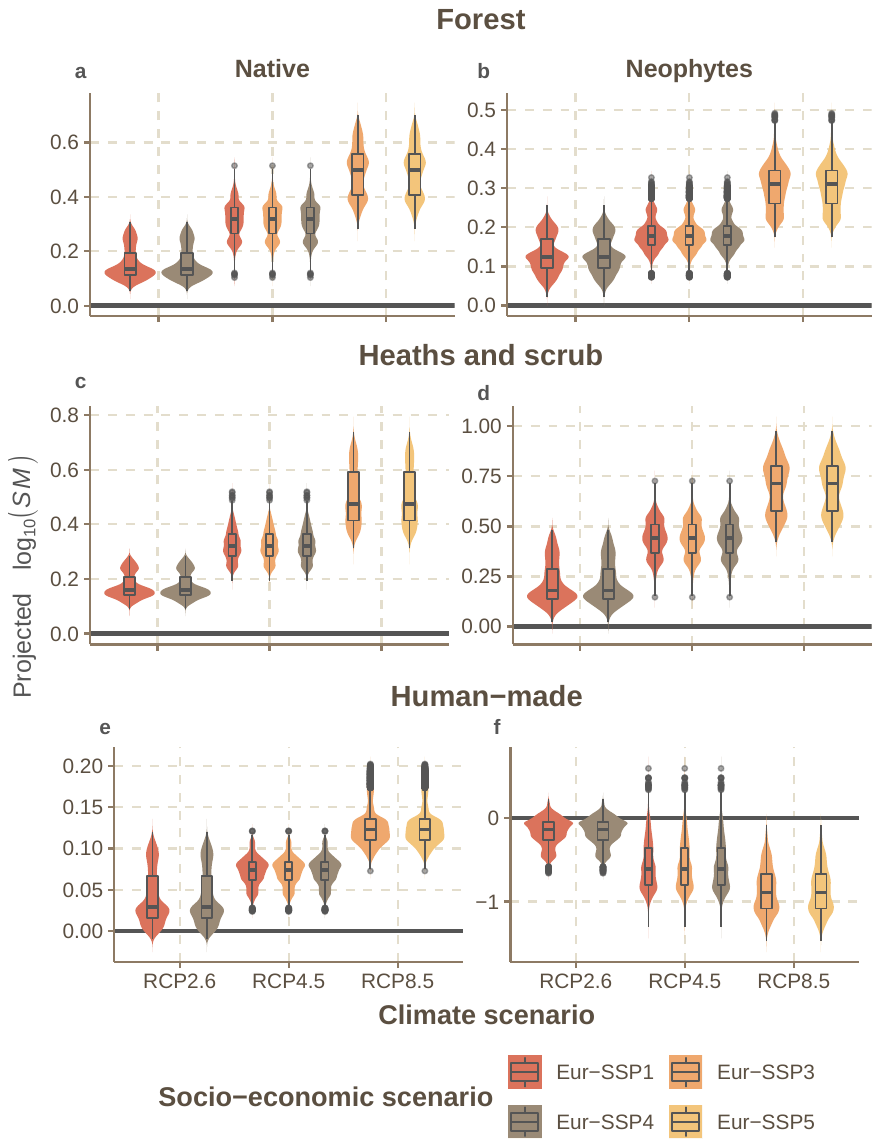

**FIGURE S7**

The projected change in the log_10_-transformed seed mass (*SM*) of *woody* plant assemblages under all plausible combined climate (RCP, all climate models pooled) and socio-economic (Eur-SSP) scenarios for 2081–2100. Other details are as in Figure S2.

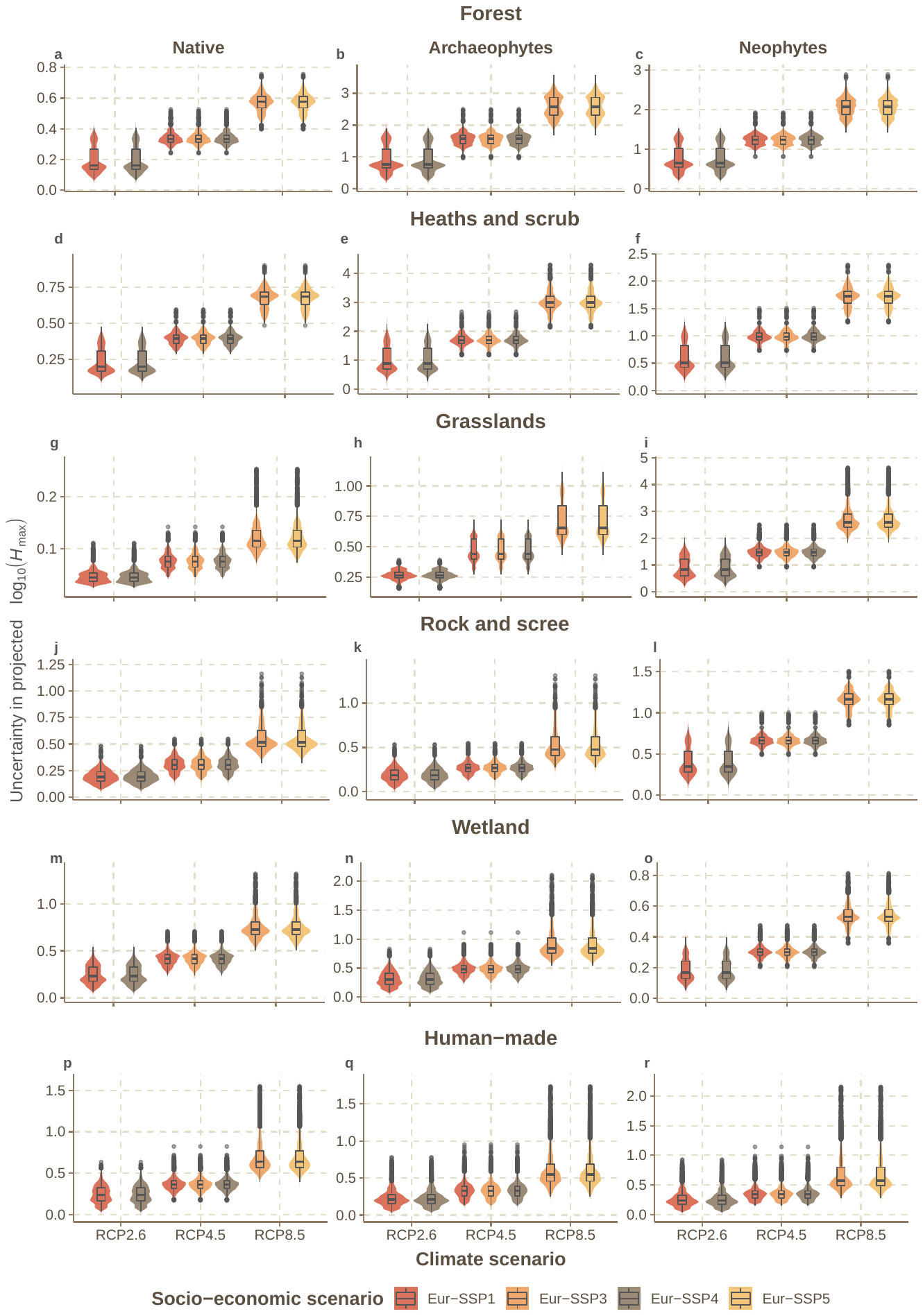

**FIGURE S8**

The uncertainty in the projected per-cell change in the log_10_-transformed maximum height (*H*_max_) of *herbaceous* plant assemblages under all plausible combined climate (RCP, all climate models pooled) and socio-economic (Eur-SSP) scenarios for 2081–2100. The uncertainty is expressed as the posterior standard deviations of per-cell model predictions.

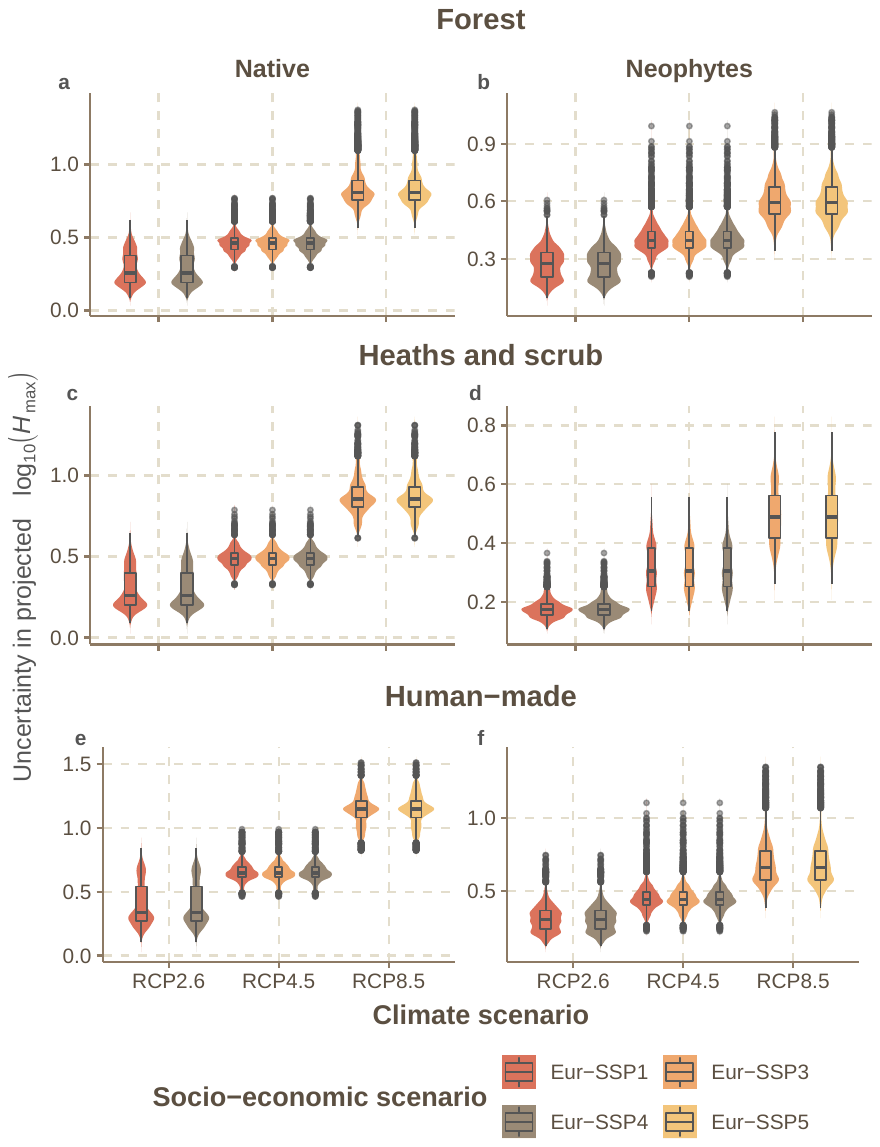

**FIGURE S9**

The uncertainty in the projected per-cell change in the log_10_-transformed maximum height (*H*_max_) of *woody* plant assemblages under all plausible combined climate (RCP, all climate models pooled) and socio-economic (Eur-SSP) scenarios for 2081–2100. The uncertainty is expressed as the posterior standard deviations of per-cell model predictions.

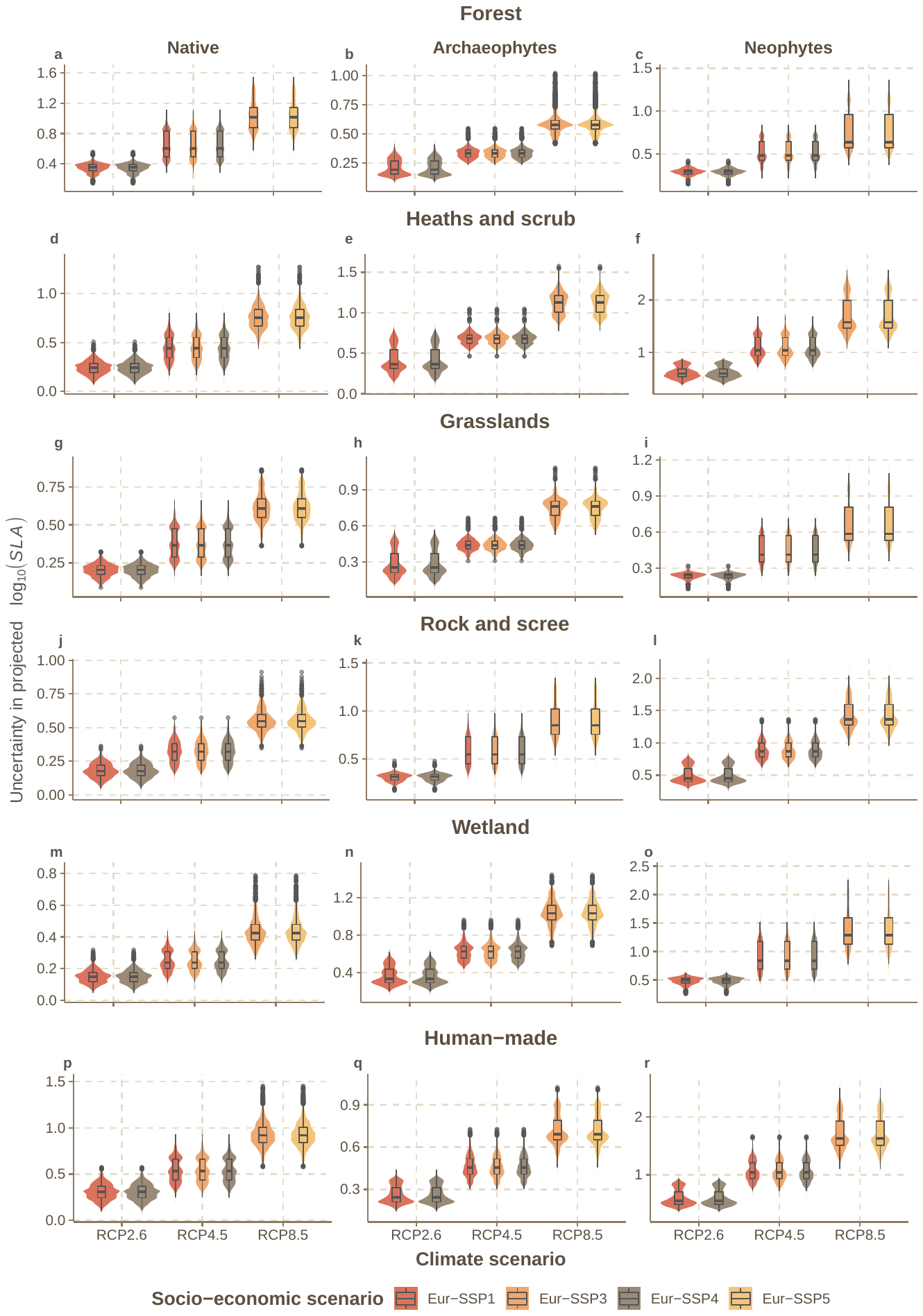

**FIGURE S10**

The uncertainty in the projected per-cell change in the log_10_-transformed specific leaf area (*SLA*) of *herbaceous* plant assemblages under all plausible combined climate (RCP, all climate models pooled) and socio-economic (Eur-SSP) scenarios for 2081–2100. The uncertainty is expressed as the posterior standard deviations of per-cell model predictions.

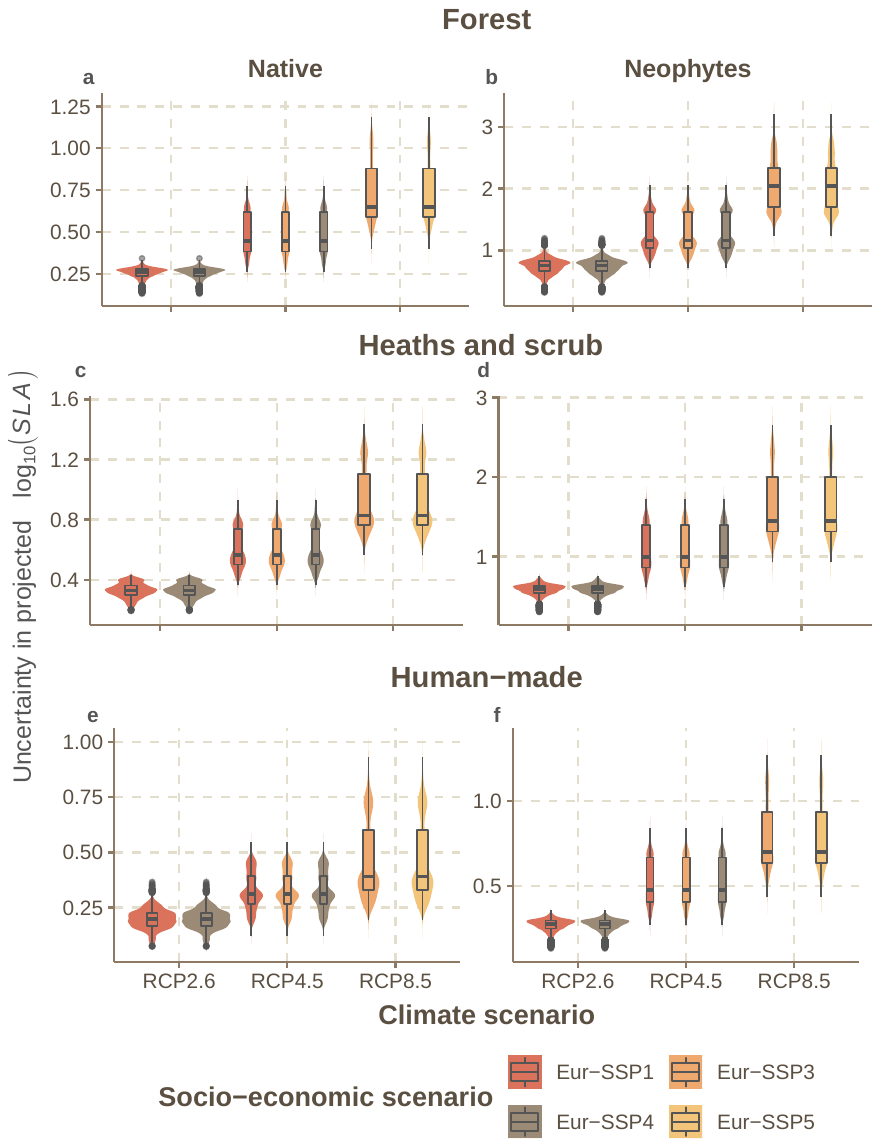

**FIGURE S11**

The uncertainty in the projected per-cell change in the log_10_-transformed specific leaf area (*SLA*) of *woody* plant assemblages under all plausible combined climate (RCP, all climate models pooled) and socio-economic (Eur-SSP) scenarios for 2081–2100. The uncertainty is expressed as the posterior standard deviations of per-cell model predictions.

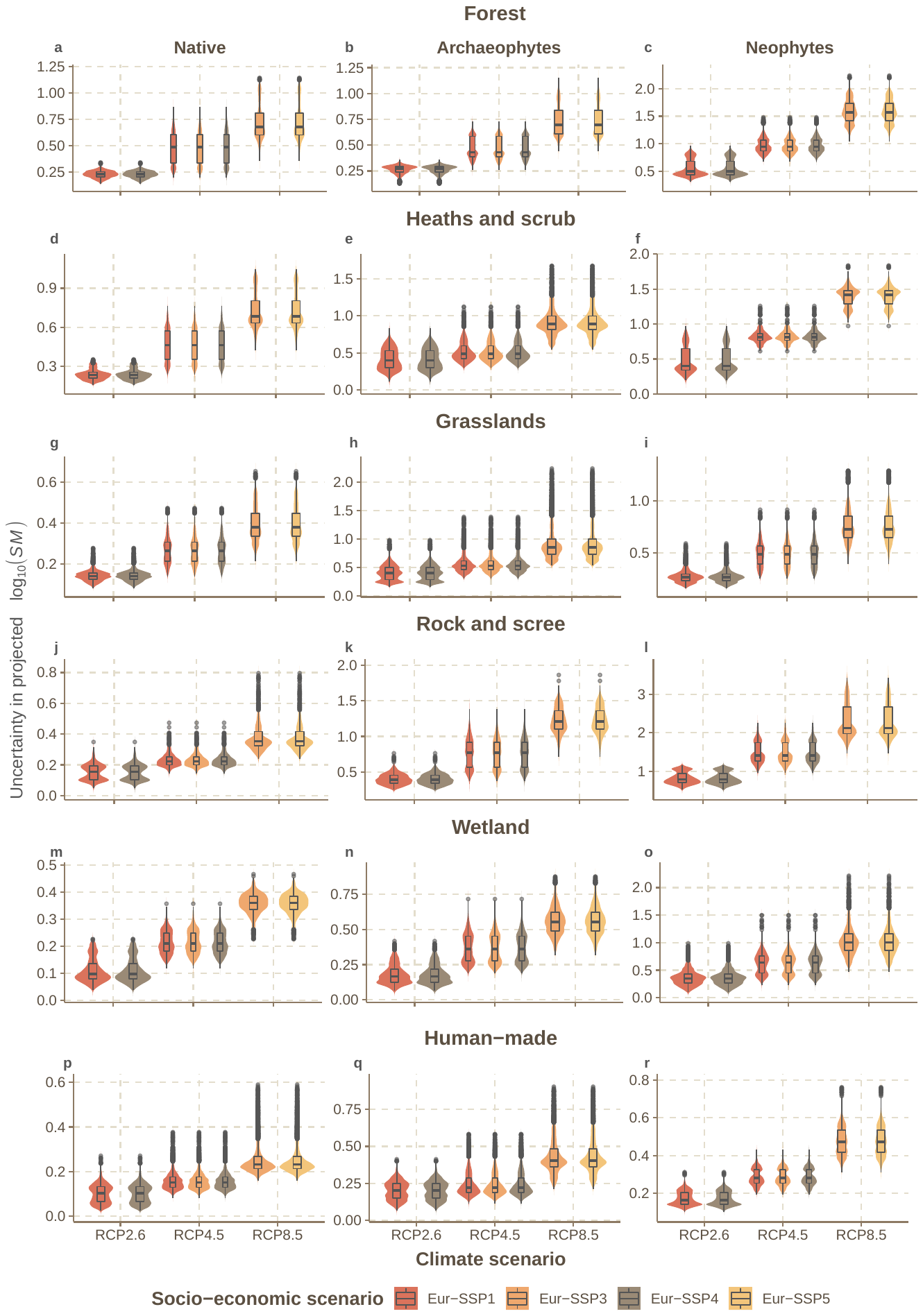

**FIGURE S12**

The uncertainty in the projected per-cell change in the log_10_-transformed seed mass (*SM*) of *herbaceous* plant assemblages under all plausible combined climate (RCP, all climate models pooled) and socio-economic (Eur-SSP) scenarios for 2081–2100. The uncertainty is expressed as the posterior standard deviations of per-cell model predictions.

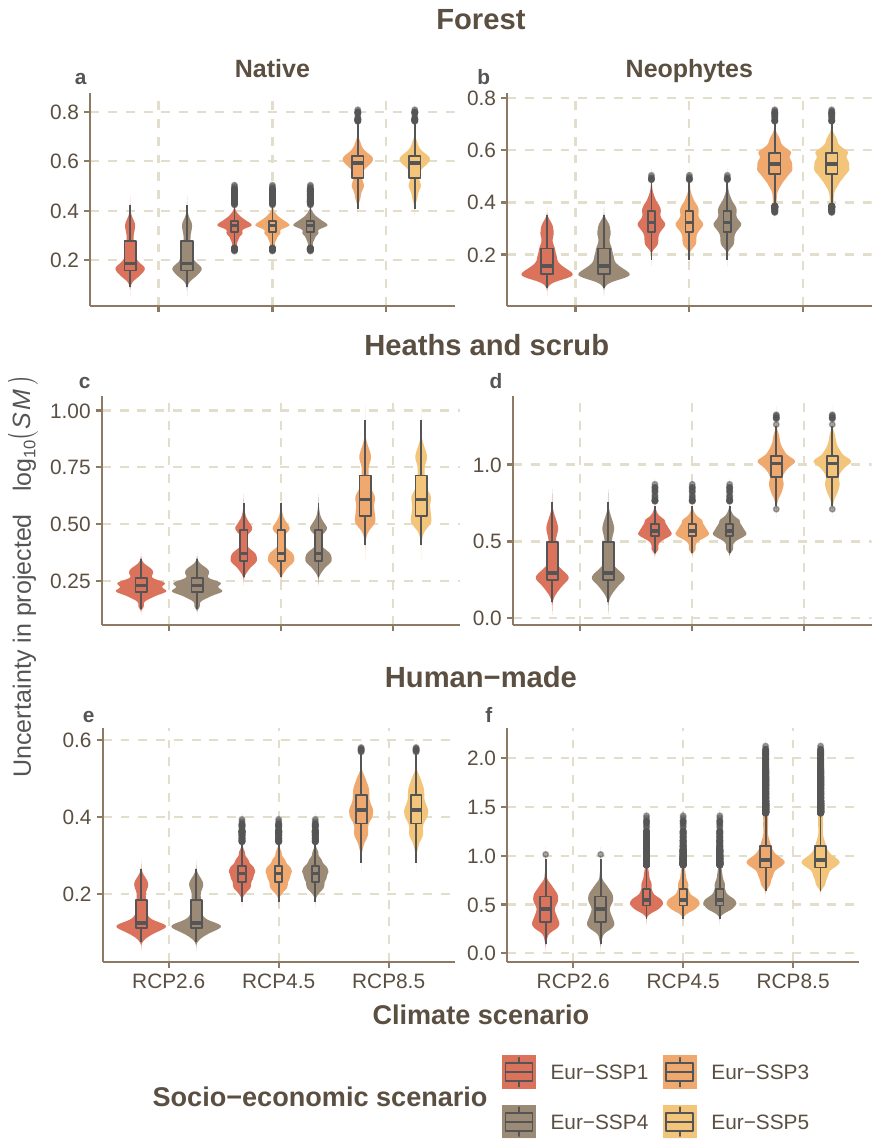

**FIGURE S13**

The uncertainty in the projected per-cell change in the log_10_-transformed seed mass (*SM*) of *woody* plant assemblages under all plausible combined climate (RCP, all climate models pooled) and socio-economic (Eur-SSP) scenarios for 2081–2100. The uncertainty is expressed as the posterior standard deviations of per-cell model predictions.

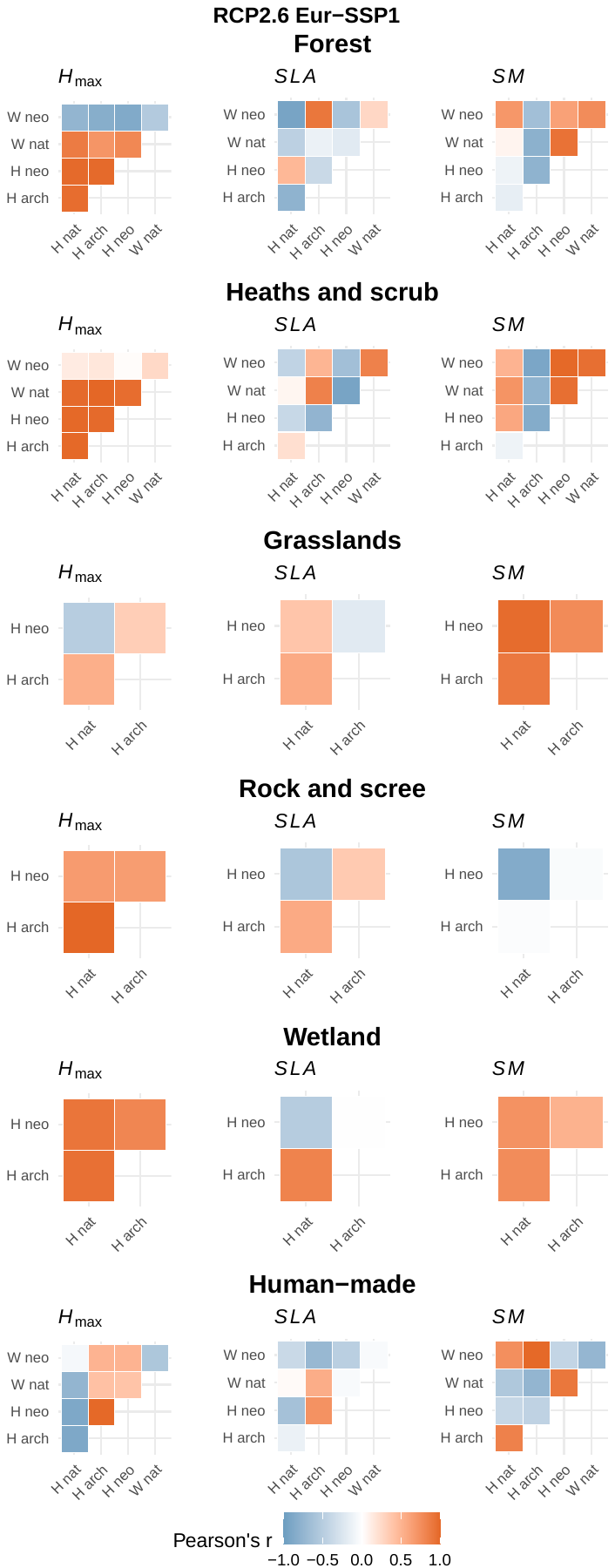

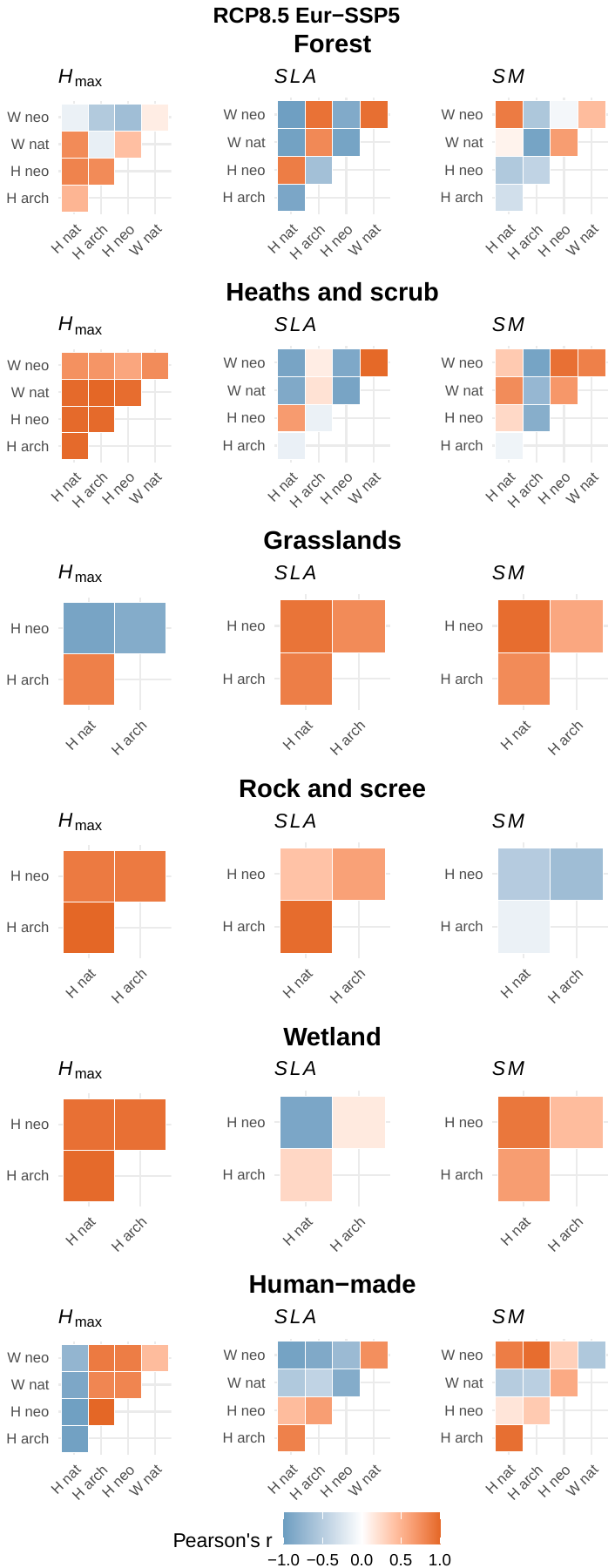

**FIGURE S14**

Correlations (Pearson’s *r*) between the projected trait change of herbaceous and woody native, archaeophyte and neophyte plant assemblages in different habitat types under RCP2.6 SSP1 and RCP8.5 SSP5 (all climate models pooled). Correlations provide insights into the extent and direction of per-cell covariation of the trait change across different assemblages. Abbreviations: *H*_max_ – maximum plant height, *SLA* – specific leaf area, *SM* – seed mass, H – herbaceous, W – woody, nat – native, arch – archaeophytes, neo – neophytes.

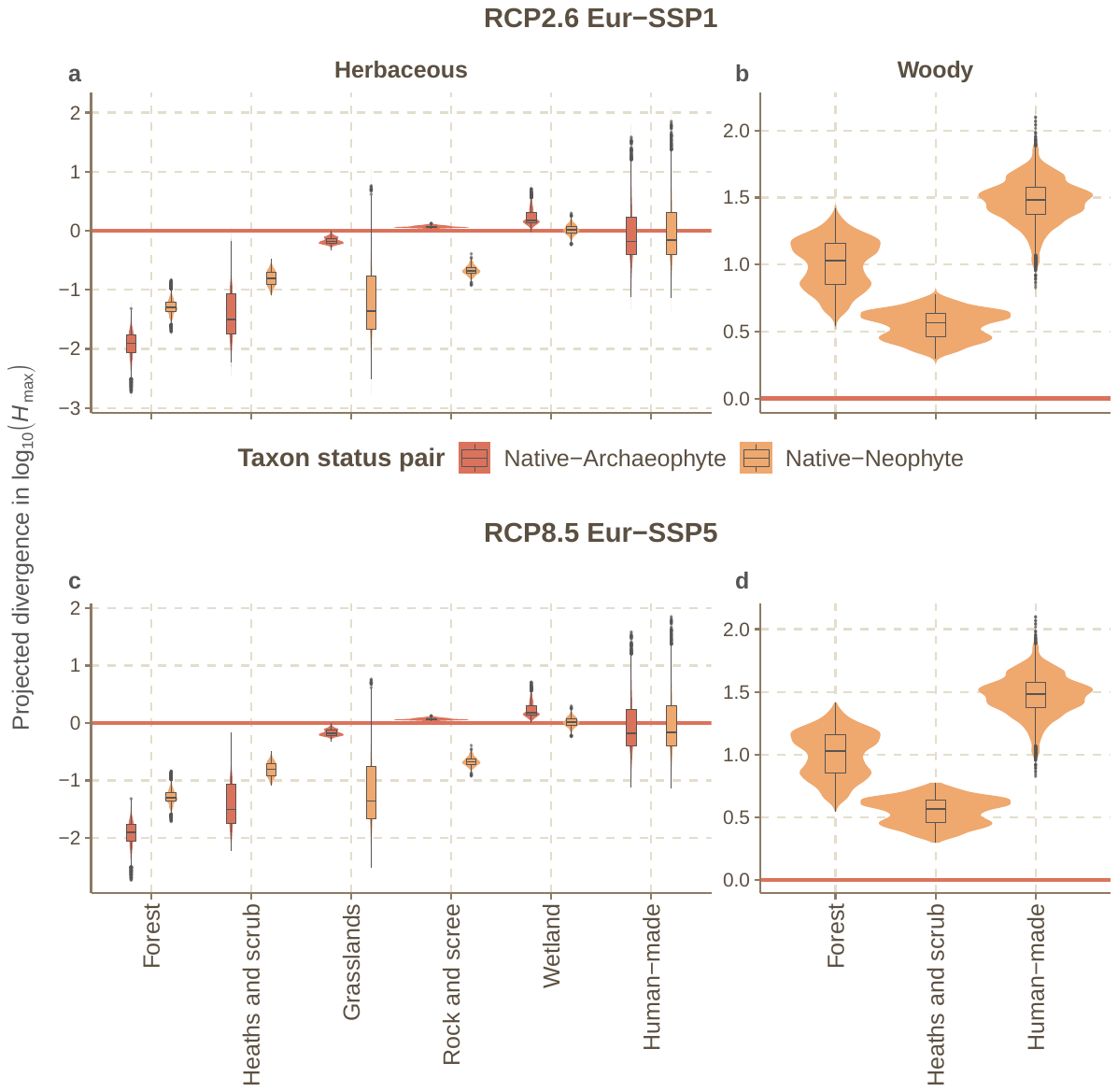

**FIGURE S15**

Difference in the projected per-cell change of maximum plant height (*H*_max_) between native and archaeophyte species and between native and neophyte species under the least extreme combined climate and socio-economic scenario (RCP2.6 Eur-SSP1, all climate models pooled) and the most extreme one (RCP8.5 Eur-SSP5) for 2081–2100. Negative values indicate that the projected *H*_max_ change for native species will be more negative compared to non-native species and positive values indicate the opposite.

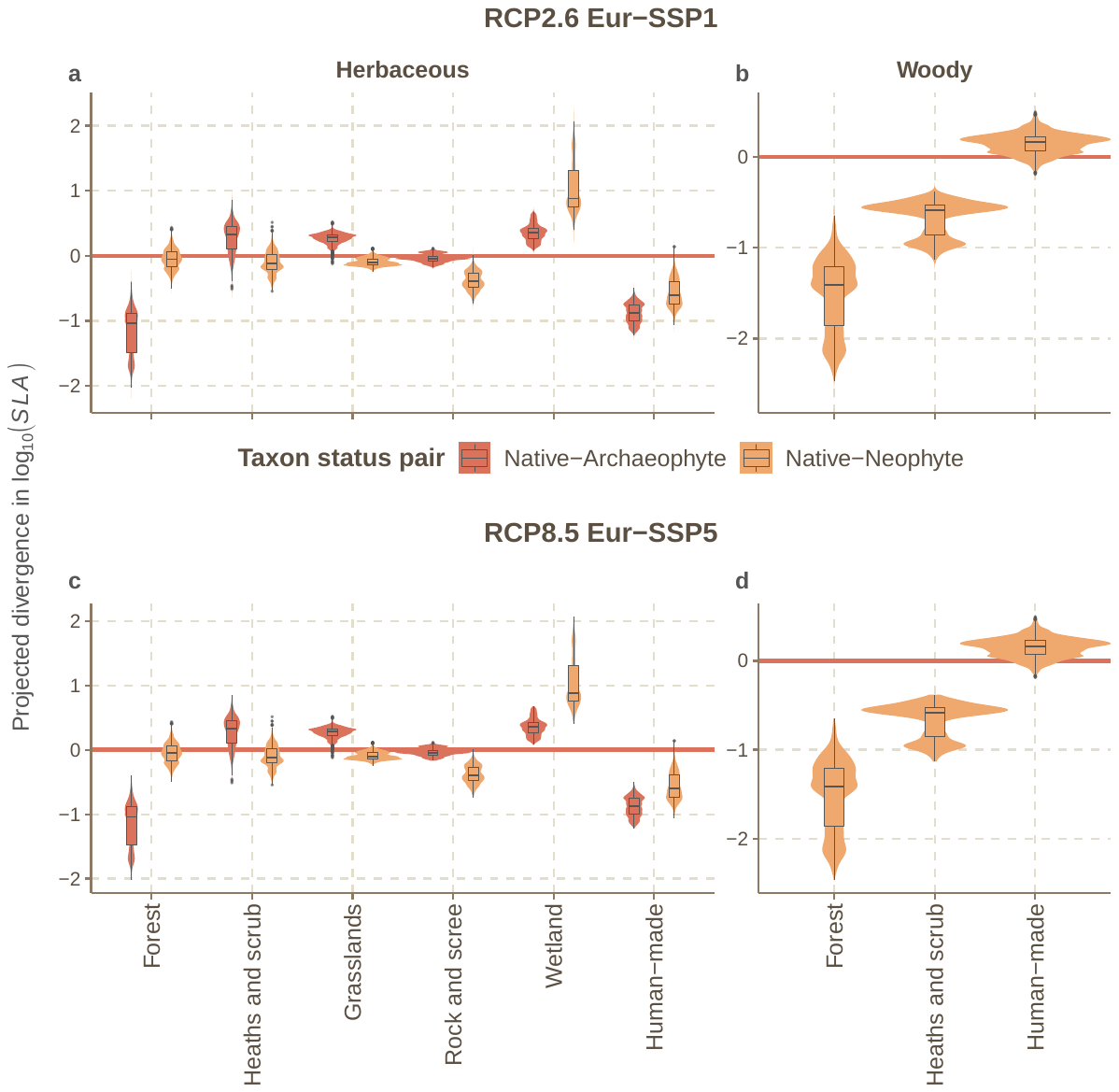

**FIGURE S16**

Difference in the projected per-cell change of specific leaf area (*SLA*) between native and archaeophyte species and between native and neophyte species under the least extreme combined climate and socio-economic scenario (RCP2.6 Eur-SSP1, all climate models pooled) and the most extreme one (RCP8.5 Eur-SSP5) for 2081–2100. Negative values indicate that the projected *SLA* change for native species is more negative compared to non-native species and positive values indicate the opposite.

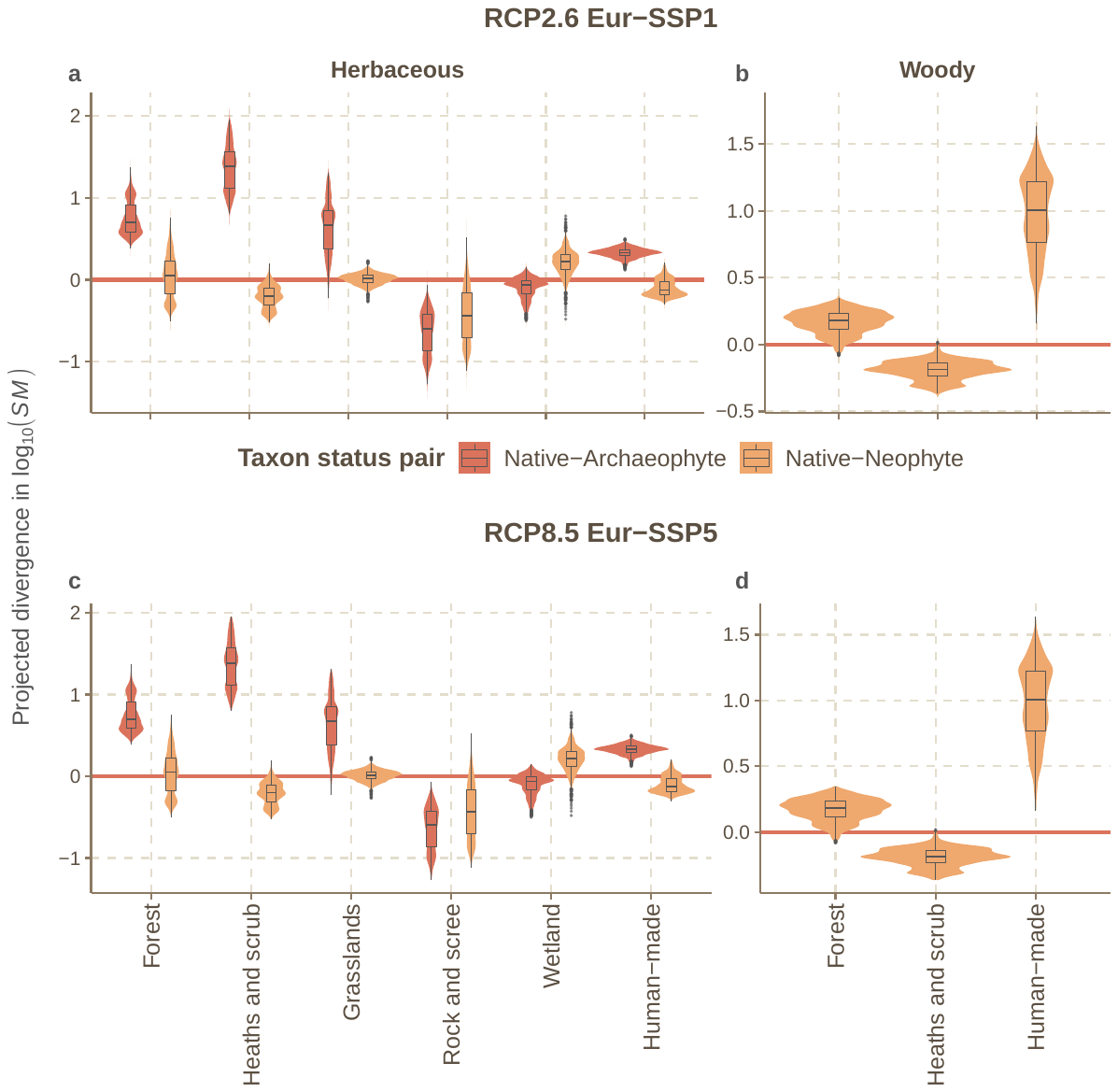

**FIGURE S17**

Difference in the projected per-cell change of seed mass (*SM*) between native and archaeophytes and between native and neophytes under the least extreme combined climate and socio-economic scenario (RCP2.6 Eur-SSP1, all climate models pooled) and the most extreme one (RCP8.5 Eur-SSP5) for 2081–2100. Negative values indicate that the projected trait change for native species will be more negative compared to non-native species and positive values indicate the opposite.

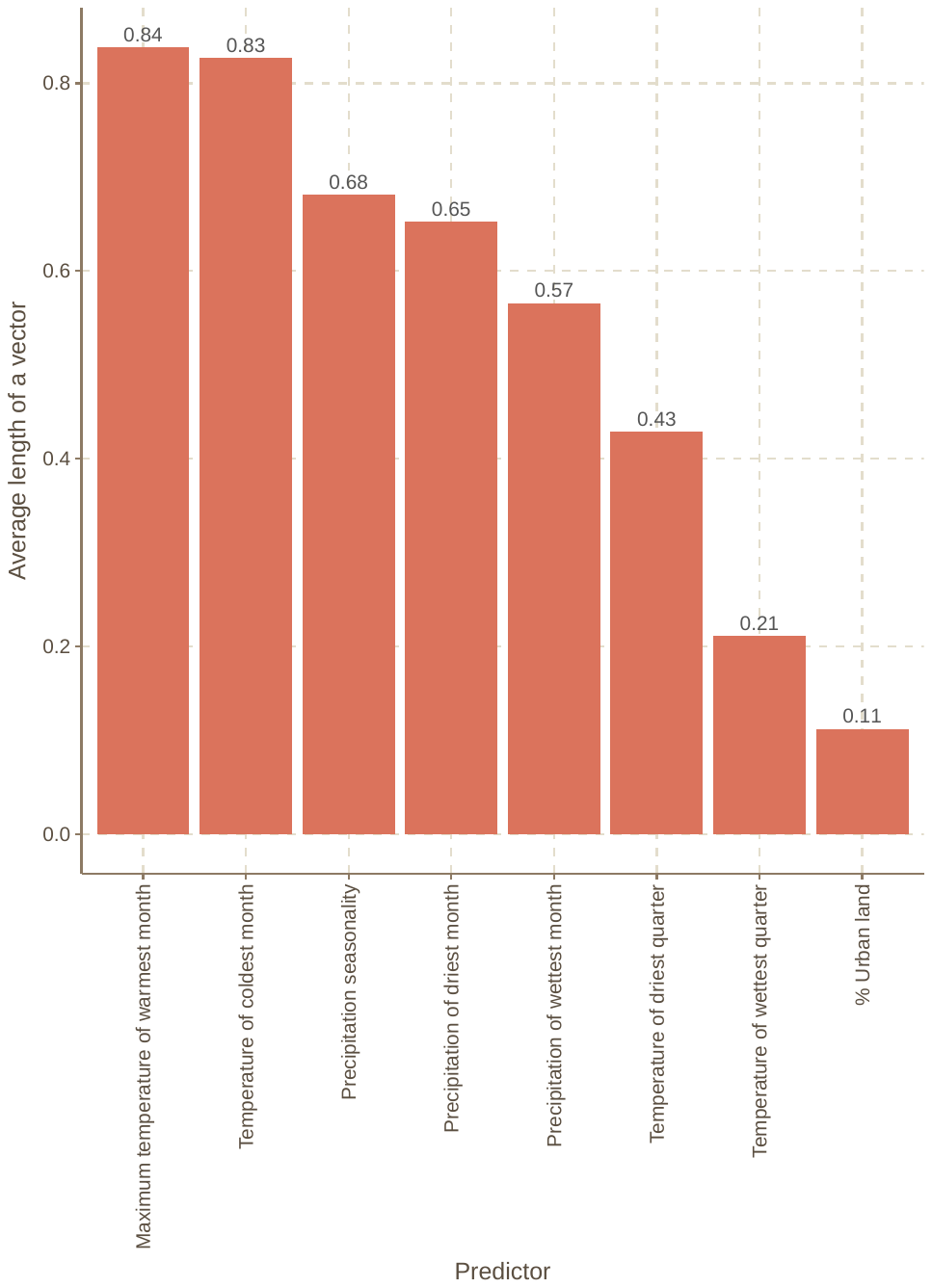

**FIGURE S18**

The overall contribution of individual predictors to the projections of the change in plant maximum height, specific leaf area, and seed mass, estimated as the average length of predictor vectors in redundancy analyses (RDA).

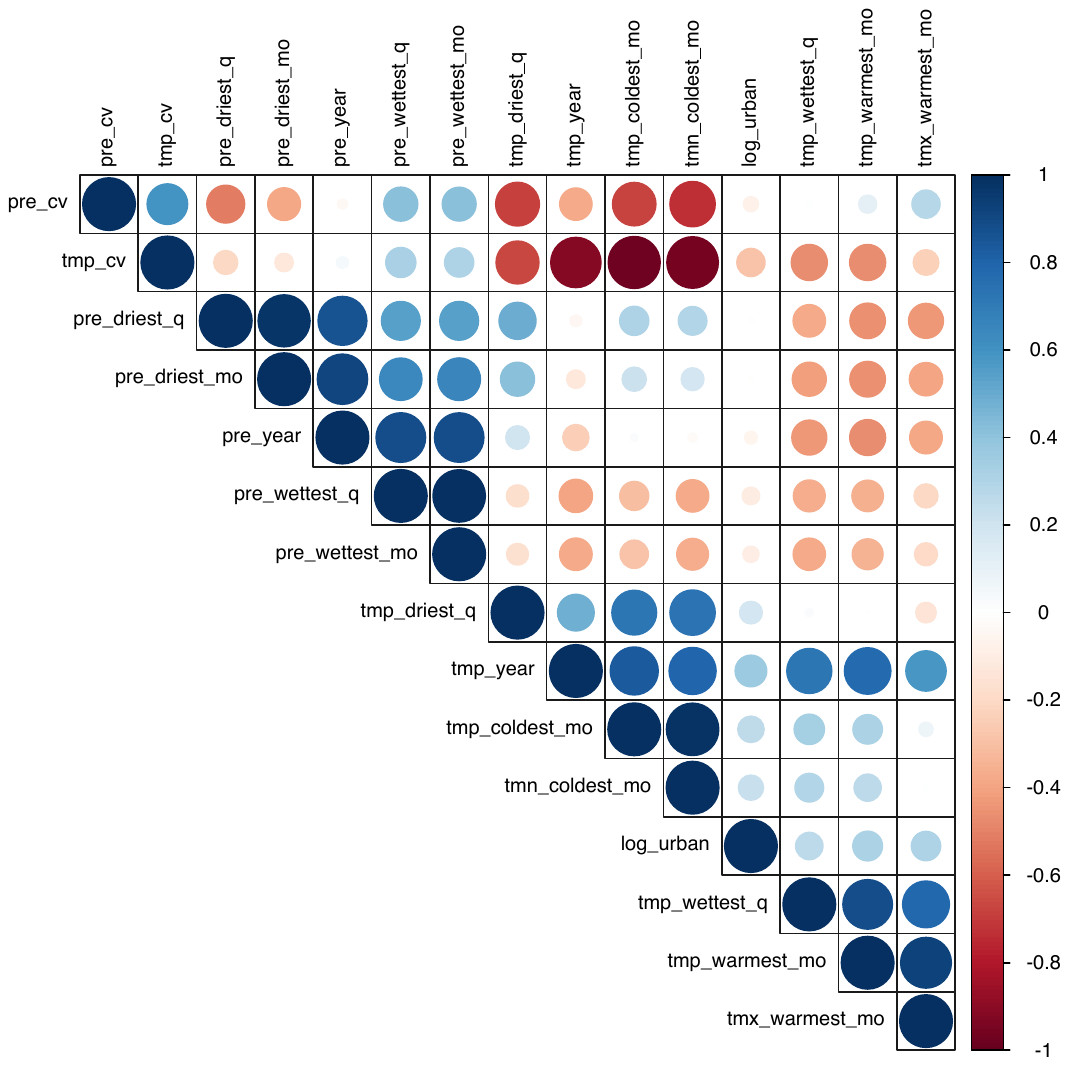

**FIGURE S19**

Pairwise correlations (Pearson’s *r*) among the environmental variables which were considered as potential predictors in the statistical models.
